# Precision of Tissue Patterning is Controlled by Dynamical Properties of Gene Regulatory Networks

**DOI:** 10.1101/721043

**Authors:** Katherine Exelby, Edgar Herrera-Delgado, Lorena Garcia Perez, Ruben Perez-Carrasco, Andreas Sagner, Vicki Metzis, Peter Sollich, James Briscoe

**Affiliations:** The Francis Crick Institute, 1 Midland Road, London, NW1 1AT, UK; Department of Mathematics, King’s College London, Strand, London WC2R 2LS, UK; Institute for Theoretical Physics, Georg-August-University Göttingen, Friedrich-Hund-Platz 1, 37077 Göttingen, Germany; Department of Mathematics, University College London, Gower Street, London WC1E 6BT, UK

## Abstract

During development, gene regulatory networks allocate cell fates by partitioning tissues into spatially organised domains of gene expression. How the sharp boundaries that delineate these gene expression patterns arise, despite the stochasticity associated with gene regulation, is poorly understood. We show, in the vertebrate neural tube, using perturbations of coding and regulatory regions, that the structure of the regulatory network contributes to boundary precision. This is achieved, not by reducing noise in individual genes, but by the configuration of the network modulating the ability of stochastic fluctuations to initiate gene expression changes. We use a computational screen to identify network properties that influence boundary precision, revealing two dynamical mechanisms by which small gene circuits attenuate the effect of noise in order to increase patterning precision. These results highlight design principles of gene regulatory networks that produce precise patterns of gene expression.

## Introduction

Embryos are characterised by remarkably organised and reproducible patterns of cellular differentiation. An illustration of this accuracy are the sharp boundaries of gene expression observed in many developing tissues. These patterns are determined by gene regulatory networks (GRNs), governed by secreted developmental signals [Davidson, 2010], raising the question of how precision is achieved inspite of the biological noise and inherent stochastic fluctuations associated with the regulation of gene expression [Raser and O’Shea, 2005].

A popular metaphor for the process of developmental pattern formation is the Waddington landscape, in which the differentiation trajectory of a cell is conceived as a ball rolling down a landscape of bifurcating valleys [Waddington, 1957]. In this representation, the landscape is shaped by the GRN with the valleys representing cell fates and developmental signals allocating cell identity by determining the valley a cell enters. This can be formalised more rigorously by describing the GRN using dynamical systems theory such that Waddingtonian valleys correspond to the attractor states of the GRN [Enver et al., 2009, Wang et al., 2011, Balázsi et al., 2011, Zhou et al., 2012]. In this view, cells can be driven out of a valley into an adjacent attractor – thus producing a change in identity – not only by developmental signals but also by gene expression noise.

How is noise buffered in developing tissues to ensure that developmental signals generate precise and reproducible patterns of gene expression? For individual genes, the activity of redundant regulatory elements (so-called shadow enhancers), the 3D architecture of the genome, the presence of multiple alleles, and the effect of RNA processing have all been proposed to buffer fluctuations and increase the robustness of gene expression [Perry et al., 2010, Frankel et al., 2010, Lagha et al., 2012, Little et al., 2013, Battich et al., 2015, Cannavò et al., 2016, Dickel et al., 2018, Osterwalder et al., 2018, Paliou et al., 2019, Tsai et al., 2019]. At the level of the tissue, mechanisms that regulate the shape, steepness or variance of gradients have been explored and their effect on the precision of gene expression detailed [Bollenbach et al., 2008, Sokolowski et al., 2012, Tkačik et al., 2015, Zagorski et al., 2017, Lucas et al., 2018]. Several mechanisms, including differential adhesion, mechanical barriers and juxtacrine signalling, have been proposed to correct anomalies and enhance precision, once cellular patterning has been initiated [Xu et al., 1999, Standley et al., 2001, Rudolf et al., 2015, Dahmann et al., 2011, Addison et al., 2018]. In addition, theoretical studies have suggested that the structure and activity of GRNs can also affect precision [Chalancon et al., 2012, Lo et al., 2015, Perez-Carrasco et al., 2016]. However, experimental evidence to support this remains elusive.

The developing vertebrate neural tube offers the opportunity to test the role of GRNs in the precision of patterning. The neural tube GRN partitions neural progenitors into discrete domains of gene expression arrayed along the dorsal-ventral axis [Sagner and Briscoe, 2019]. The boundaries between these domains are clearly delineated and accurately positioned [Kicheva et al., 2014] resulting in sharp spatial transitions in gene expression that produce characteristic stripes of molecularly distinct cells. In the ventral neural tube, the secreted ligand Sonic Hedgehog (Shh), emanating from the notochord and floor plate, located at the ventral pole, controls the pattern forming GRN (Fig. 1A). The regulatory interactions between the transcription factors (TFs) comprising the GRN explain the dynamics of gene expression in the ventral neural tube and produce the genetic toggle switches that result in discrete transitions in gene expression in individual cells [Balaskas et al., 2012]. The network includes the TFs Pax6, Olig2, Irx3 and Nkx2.2. Irx3 represses Olig2 [Novitch et al., 2001], while Nkx2.2 is repressed by Pax6, Olig2 and Irx3 [Briscoe et al., 1999, Briscoe et al., 2000, Novitch et al., 2001, Balaskas et al., 2012]. In the absence of Shh signaling, progenitors express Pax6 and Irx3. Moderate levels of Shh signalling are sufficient to induce Olig2 expression and repress Irx3 to specify motor neuron progenitors (pMN) [Ericson et al., 1997, Briscoe et al., 2000, Novitch et al., 2001, Balaskas et al., 2012]. In response to high and sustained levels of Shh signalling, Nkx2.2 is induced and inhibits the expression of Pax6 and Olig2, which then generates p3 progenitors and delineates the p3-pMN boundary (Fig. 1B). In embryos lacking Pax6, the domain of Nkx2.2 expression expands resulting in a decrease in Olig2 expression and dorsal shift in the p3-pMN boundary [Ericson et al., 1997, Novitch et al., 2001, Balaskas et al., 2012].

**Figure 1:**
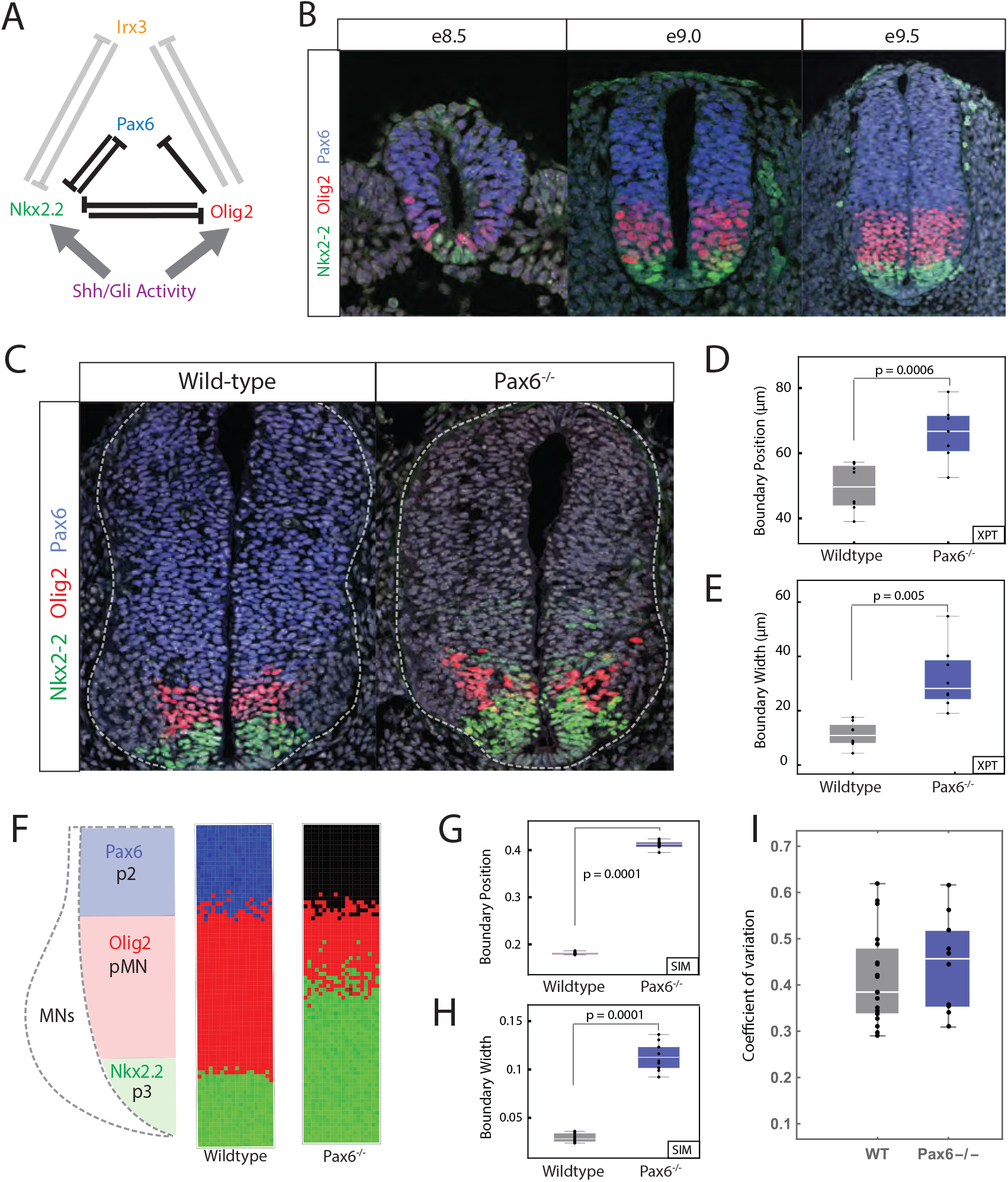
Pax6 contributes to boundary precision. (**A**) Schemaic of the GRN responsible for positioning the p3 and pMN domains. (**B**) Immunofluorescence assays of Pax6 (blue), Olig2 (red) and Nkx2.2 (green) in neural progenitors from e8.5 to e9.5. (**C**) WT and Pax6^−/−^ embryos assayed for Olig2, Pax6 and Nkx2.2. (**D**) Position of the pMN-p3 boundary in WT (grey) and Pax6^−/−^ (blue). (Box plots in all figures show upper and lower quartile and mean; *n* = 7 (WT), *n* = 8 (Pax6^−/−^), Mann-Whitney test *p* = 0.005). (**E**) Width of pMN-p3 boundary in WT (grey) and Pax6^−/−^ (blue) (Mann-Whitney test *p* = 0.0006). (**F**) Stochastic simulations of the GRN in WT (middle) and Pax6^−/−^ (right). (**G,H**) Boundary position and width from simulations. Width is given as fraction of total neural tube size. *n* = 10 (WT), *n* = 10 (Pax6^−/−^), Mann-Whitney test *p* = 0.0001 for position and boundary width. (**I**) Coefficient of variation (CV) of Olig2 levels for WT and Pax6^−/−^ (Mann-Whitney *p* = 0.422)

In addition to the change in the position of the p3-pMN boundary, the loss of Pax6 also results in decreased precision of the p3-pMN boundary with noticeably more intermixing of cells [Ericson et al., 1997, Briscoe et al., 2000, Novitch et al., 2001, Balaskas et al., 2012]. Here we set out to understand what explains this loss of precision. We hypothesised that stochastic fluctuations in gene expression coupled with changes in the dynamics of the GRN in the absence of Pax6 account for the decreased boundary precision. We provide a combination of experiment, data analysis and theory that are consistent with this idea. We also found that perturbing the regulatory input onto Olig2, by deleting a single cis-regulatory element, altered the dynamics of the GRN and decreased the precision of the p3-pMN boundary. The decreased precision was not a result of increased noise in the expression of individual genes. Instead the absence of the Olig2 regulatory element, similar to the loss of Pax6, changed the overall configuration of the stochastic fluctuations and made transitions from a pMN to p3 state more likely. A computational screen for networks that generate precise boundaries supported this idea and revealed two dynamical mechanisms by which small gene circuits attenuate the effect of noise in order to increase patterning precision. Thus, although mechanisms necessitating additional signals, differential adhesion or cell mechanics are often invoked to explain the precision of tissue patterning, our analysis suggests that the intrinsic properties of a GRN can also enhance boundary precision.

## Results

### Pax6 contributes to p3-pMN boundary precision

We assayed the precision of the boundary between p3 (Nkx2.2 expressing) and pMN (Olig2 and Pax6 expressing) in the ventral neural tube. Consistent with previous reports [Ericson et al., 1997, Balaskas et al., 2012], compared to wild-type (WT) mouse embryos, the precision of the boundary between p3-pMN domains was decreased in embryos lacking Pax6, resulting in more intermixing of cells expressing Olig2 or Nkx2.2 (Fig. 1C) [Ericson et al., 1997, Briscoe et al., 2000, Balaskas et al., 2012]. Measurements of the dorsal-ventral width of the region that contains both Nkx2.2 and Olig2 expressing cells in WT and Pax6 mutant embryos (Supplemental Section G.8) indicated that between e9.0 and e10.5, the width of the pMN-p3 boundary region is wider in Pax6^−/−^ embryos, consistent with a loss of precision (Fig. 1E & S1).

We hypothesised that the decreased precision of the Nkx2.2 boundary, leading to the increased width in Pax6^−/−^ embryos (Fig. 1C), could be explained by noise in gene expression in the GRN. Previously, we established a deterministic model of the GRN, based on coupled Ordinary Differential Equations (ODEs), that replicated the response of the network to Shh signalling and the shifts in boundary position in mutant embryos, including Pax6^−/−^ [Panovska-Griffiths et al., 2013, Balaskas et al., 2012, Cohen et al., 2014]. This model recapitulated the pMN and p3 steady states. The analysis also produced a region of bistablity in which both the pMN and p3 states were stable, however, due to the initial conditions and deterministic behaviour, the system always adopted a pMN state in the bistable region. We reasoned that fluctuations in gene expression could result in noise driven transitions within the bistable region from a pMN state to a p3 identity. (For a glossary of dynamical systems terminology see Supplemental Section B.) We constructed a stochastic differential equation (SDE) model that retained the parameters of the ODE model but incorporated a description of intrinsic gene expression fluctuations, based on experimental measurements (Supplemental Section C & D). Simulations with this model revealed that stochasticity in gene expression was sufficient to promote a switch from a pMN state to a p3 identity within the bistable region and the probability of a noise driven transition increased with higher levels of signal as the system approached the p3 monostable regime. Moreover, the hysteresis that is a consequence of the bistablity [Panovska-Griffiths et al., 2013, Balaskas et al., 2012, Cohen et al., 2014] meant that transitions from pMN to p3 were more frequent than the reverse.

We used the SDE model to simulate a Pax6^−/−^ mutant. Compared to WT simulations, in the Pax6^−/−^ mutant not only was the boundary of the Nkx2.2 expressing p3 domain displaced dorsally, but the boundary also showed markedly reduced precision (Fig. 1F,G,H). Thus, inclusion of intrinsic noise in the GRN dynamics was sufficient to accurately reproduce the alterations in the position and precision of gene expression boundaries.

### An Olig2 enhancer influences boundary precision

To test the hypothesis that the regulatory dynamics of the GRN affect the the precision of patterning we sought to alter the strength of interactions within the network. We turned our attention to the cis-regulatory elements controlling the TFs in the GRN, as regulatory elements have been shown to affect the reliability of patterning in other systems [Perry et al., 2011, El-Sherif and Levine, 2016]. Several predicted regulatory regions are located in the vicinity of Olig2; these include a prominent candidate region 33kb upstream of the Olig2 gene [Oosterveen et al., 2012, Peterson et al., 2012], which we termed O2e33. This region binds (i) the repressor Nkx2.2; (ii) Sox2, which activates Olig2; and (iii) the Gli proteins, the transcriptional effectors of the Shh pathway (Fig. 2A) [Oosterveen et al., 2012, Peterson et al., 2012, Nishi et al., 2015, Kutejova et al., 2016] and becomes accessible in neural progenitors [Metzis et al., 2018]. To test the role of O2e33 in the network we first analysed its function *in vitro* in neural progenitors differentiated from mouse embryonic stem (ES) cells [Gouti et al., 2014]. Unlike WT cells, which express high levels of Olig2 at Day 5 of differentiation [Gouti et al., 2014, Sagner et al., 2018], cells in which the O2e33 enhancer had been deleted (O2e33^−/−^) had a marked reduction in levels of Olig2. By Day 6, Olig2 expression had increased in O2e33^−/−^ cells, but the percentage of cells and the level of expression never reached that of WT (Fig. 2B,C). Consistent with the role of Olig2 in the generation of MNs, the production of MNs was substantially decreased in O2e33^−/−^ cells (Fig. 2D).

**Figure 2:**
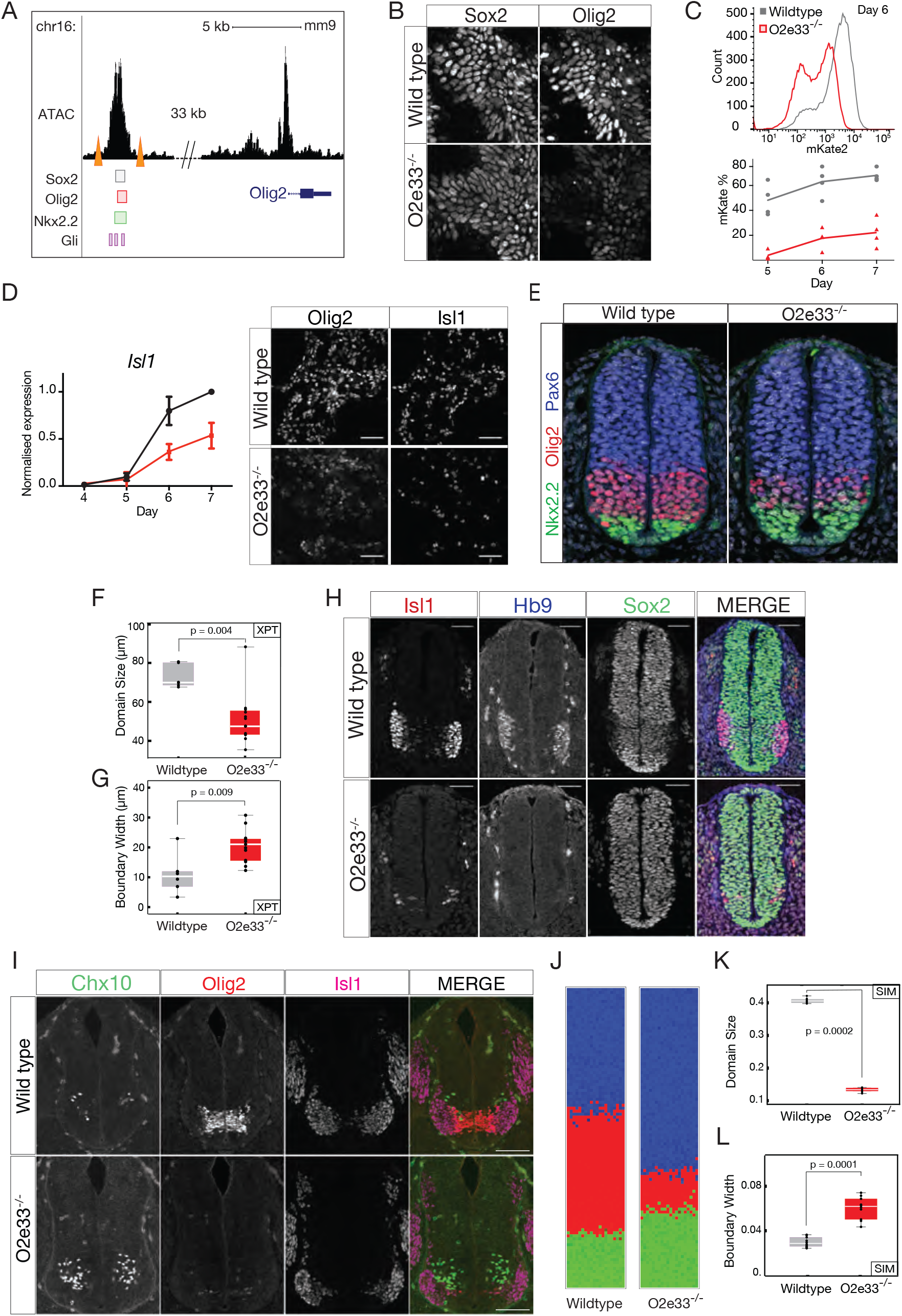
An Olig2 enhancer affects precision of the pMN-p3 boundary. (**A**) Chromatin accessibility (ATAC-seq) and predicted TF binding locations around Olig2. CRISPR target sites (orange triangles) for deletion of the O2e33^−/−^ [Metzis et al., 2018, Kutejova et al., 2016, Peterson et al., 2012, Oosterveen et al., 2012]. (**B**) Sox2 (expressed in all neural progenitors) and Olig2 at day 6 in neural progenitors differentiated from WT and O2e33^−/−^ ES cells exposed to 500nM SAG. (**C**) Flow cytometry (top) for mKate2 flourescence in Olig2-T2A-mKate2 ES cell derived neural progenitors exposed to 500nM SAG. (**D**) RT-qPCR indiates Isl1 is decreased in O2e33^−/−^ (red) cells compared to WT (black) cells differentiated under spinal cord conditions. Similarly, Olig2 and Isl1 expressing cells are reduced in mutant compared to WT. (**E**) Olig2, Pax6 and Nkx2.2 in transverse sections of e9.5 neural tube from WT and O2e33^−/−^ (red, Olig2; green, Nkx2.2). (**F, G**) Domain size and boundary width in WT (grey) and O2e33^−/−^ mutants (red). *n* = 6 (WT), *n* = 12 (O2e33), Mann-Whitney test *p* = 0.004. The p3-pMN boundary is wider in O2e33^−/−^ mutants compared to WT (Mann-Whitney test *p* = 0.009). (**H**) Isl1 and Hb9 expressing motor neurons are reduced in O2e33^−/−^ embryos compared to WT. (**I**) Chx10 expressing V2 neurons increase in the 02e33^−/−^ mutant. Scale bars = 100*μ*m. (**J**) Simulations of the O2e33^−/−^ model recapitulate *in vivo* observations of a narrower pMN domain and decreased precision of the p3-pMN boundary. (**K, L**) Boundary width (I) and position (H) from simulations (box plot shows upper and lower quartile and mean; *n* = 10 (WT); *n* = 10 (O2e33), Mann-Whitney test *p* = 0.0001 for both).

We used the experimentally observed delay in Olig2 induction to identify changes in model parameters that mimic the effect of deleting the O2e33 enhancer (Supplemental Section E). Of the parameter sets that delayed Olig2 induction *in silico*, most predicted the generation of a smaller pMN domain, resulting from a ventral shift in the dorsal boundary. Strikingly, many of the parameter sets also predicted a loss of boundary sharpness of the p3-pMN boundary. To test these predictions, we generated mutant mice lacking the O2e33 enhancer (Methods). Assaying the neural tube of embryos from these mice revealed lower Olig2 expression levels in pMN cells and a delay in the induction of Olig2 in O2e33^−/−^ embryos compared to WT, in agreement with *in vitro* results (Fig. S2, S3). As predicted by the *in silico* analysis, the pMN domain was decreased in size in O2e33^−/−^ embryos, with its dorsal limit of expression noticeably more ventrally positioned (Fig. 2E). Moreover, the boundary between the pMN and p3 domain was less precise than WT (Fig. 2E, F,G). Consistent with the reduced domain size, there was a significant reduction in the generation of MNs in O2e33^−/−^ embryos and a comcommitant increase in V2 neuron production (Fig. 2H,I). The decrease in the precision of the boundary, despite continued expression of Olig2 and Pax6 in pMN cells, suggests that secondary correction mechanisms do not suffice to determine the precision of the boundary between these two domains.

Using the *in vivo* observations we further constrained the parameter space of the model by restricting our analysis to parameter sets that generated an imprecise p3-pMN boundary and alter the position of the pMN-p2 boundary (Supplemental Section E). This produced simulations in which the loss of boundary precision in the O2e33^−/−^ embryos is not as severe as the Pax6^−/−^ phenotype, in line with the experimental data (Fig. 2J), and the width of the boundary and the size of the pMN domain were consistent with *in vivo* analysis (Fig. 2K-L). Taken together, the data suggest that Pax6 function and the activity of the O2e33 enhancer increase the precision of the p3-pMN boundary by attenuating the effects of gene expression stochasticity in the GRN.

### Rate of stochastic switching is controlled by GRN dynamics

To understand the mechanism by which Pax6 and O2e33 contribute to boundary precision, we explored the dynamical properties of the SDE model. The model did not predict a difference in the magnitude of the fluctuations in the expression of individual genes between the WT and the Pax6 mutant (Supplemental Section C). Consistent with this, experimental measurements of the coefficient of variation (CV) of Olig2 from WT and Pax6^−/−^ embryos did not reveal significant differences (Fig. 1I). This raised the possibility that, rather than the size of fluctuations in individual genes, the change in precision was a consequence of the regulatory interactions of the network. The model of the GRN predicts a bistable regime between the two steady states of Nkx2.2 (p3) and Olig2/Pax6 (pMN) (Fig. 3A) [Balaskas et al., 2012, Cohen et al., 2014]. In the absence of noise, the transition between the two steady states is determined solely by the level of Shh signalling. However, in the presence of intrinsic noise, fluctuations in gene expression can result in spontaneous transitions between pMN and p3 identity within the bistable region [Perez-Carrasco et al., 2016]. Below a threshold of Shh signalling, the rate of transitions is very low and cells remain in the pMN state. Conversely, above a certain level of Shh signalling, transitions from the pMN to the p3 steady state take place so rapidly that almost all cells undergo this transition. In between these two regimes, a region of heterogeneity is observed in which there is an intermediate probability for each cell to spontaneous transition (≤50 hours), see Fig. 3A-B. We calculated the characteristic time it would take for transitions between the pMN and p3 states at different dorsal-ventral positions of the neural tube. We termed this “fate jump time”. For WT, fate jump time changes rapidly in response to Shh signalling, implying that there is only a limited region where the effective probability of transitions is not 0 or 1 (Fig. 3B; black line). By contrast, the larger region of heterogeneity observed in the Pax6^−/−^ mutant is due to the weaker dependence of fate jump time on levels of Shh signalling (Fig. 3B; blue line). There is a larger range of Shh levels for which noise driven transitions are possible and therefore a larger boundary region where cells in both p3 and pMN states exist.

**Figure 3:**
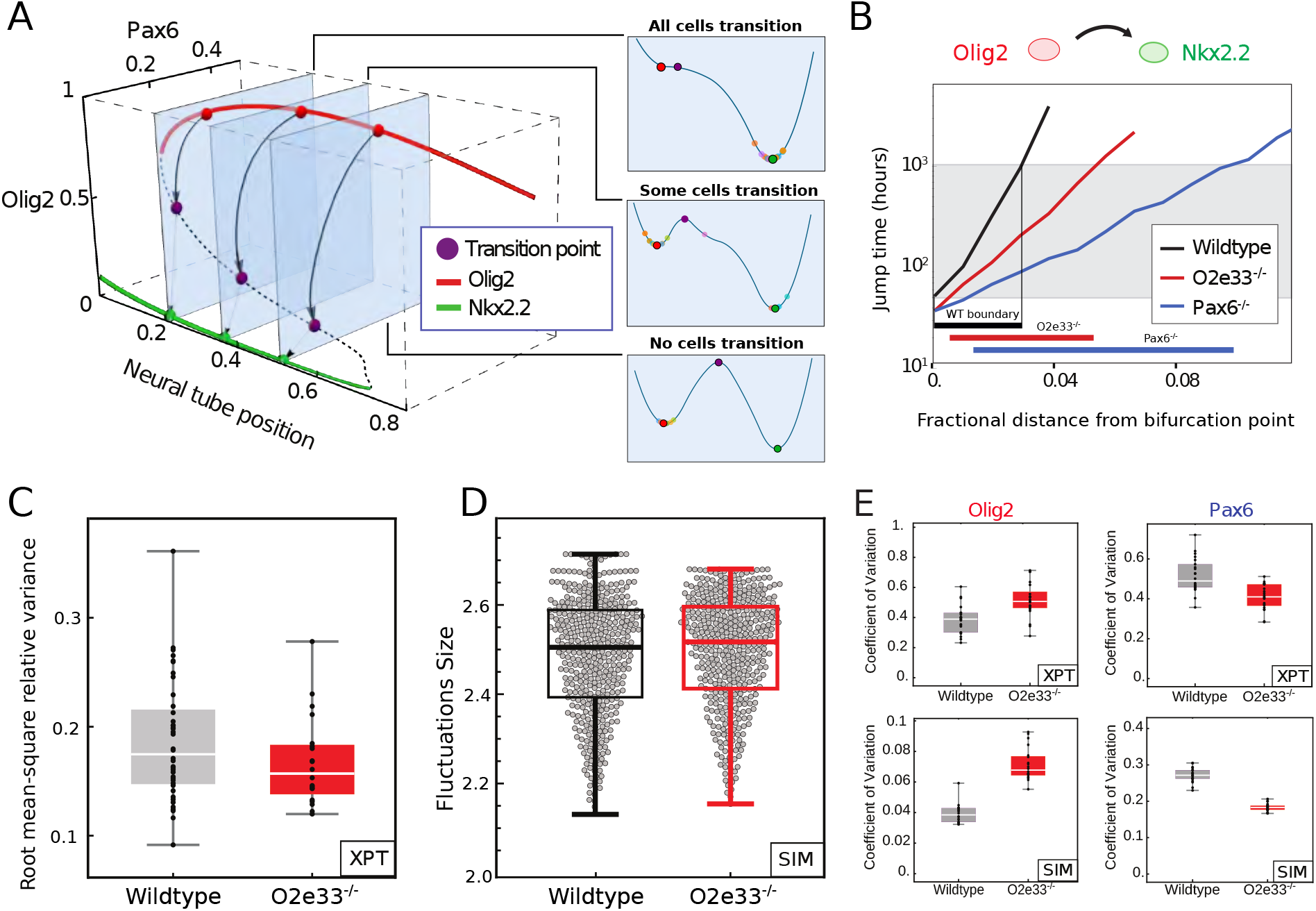
The rate of transition between progenitor states is determined by the GRN structure. (**A**) A 3D bifurcation diagram illustrates bistablity for pMN (red; expressing Olig2 and Pax6) and p3 (green; expressing Nkx2.2) with a transition point (unstable fixed point of dynamics, purple). Noise driven transition pathway from pMN to p3 is indicated by black arrows. Panels (right) represent the transitions as one-dimensional Waddington landscape sketches. (**B**) Fate jump times calculated from simulations: pMN to p3 in WT (black), Pax6^−/−^ (blue) and O2e33^−/−^ mutants (red). Fractional distance refers to distance from the bifurcation point. Grey shading indicates where transitions can occur on developmental timescales. (**C**) Total variance in gene expression per embryo (Olig2 and Pax6) within pMN domain for WT (grey) and O2e33^−/−^ embryos (red). Relative root-mean-square variance of WT and O2e33^−/−^ embryos captures total noise of the system. No significant change in noise levels between genotypes (*p* > 0.05, Mann-Whitney test). (**D**) Measurements of noise *in silico* in the pMN domain in WT and O2e33^−/−^, each grey point is an individual configuration (Supplemental Section C, Mann-Whitney test *p* > 0.05). (**E**) Coefficient of variation for Olig2 (left) and Pax6 (right) in WT (grey) and O2e33^−/−^ (red) from experimental data (top) and *in silico* simulations (bottom).

Fate jump times changed more slowly for O2e33^−/−^ than for WT (Fig. 3B), but more rapidly than for the Pax6^−/−^ system. This is in line with the boundary precision of O2e33^−/−^ embryos falling between that of WT and Pax6^−/−^. Analysis *in vivo* of the magnitude of the combined fluctuations in Pax6 and Olig2 indicated that it was similar in WT and O2e33^−/−^ (Fig. 3C; Supplemental Section G.8). Consistent with this, the combined magnitude of fluctuations of Pax6 and Olig2 in simulations were similar in WT and O2e33^−/−^ mutants. This suggested that, similar to Pax6^−/−^ embryos, the decreased precision was not the result of an increase in the overall magnitude of fluctuations (Fig. S9)(Fig. 3D). In addition, simulations of the O2e33^−/−^ mutant predicted that variability in Olig2 should increase whereas the variability of Pax6 should decrease. In agreement with this, the CV of Olig2 and Pax6 between WT and O2e33^−/−^ *in vivo* were increased and decreased, respectively (Fig. 3E).

### The dynamical landscape determines boundary precision

To investigate the reasons for the change in fate jump time in O2e33^−/−^ and Pax6^−/−^ mutants, we explored the effect of these perturbations on the dynamical landscape of the system (see Supplemental Section B). Transitions between p3 and pMN states involve the system passing through, or very close to, a point in gene expression space - the saddle point in the dynamical landscape - that is characterised by specific levels of the transcription factors (TFs), we refer to this as the “transition point” (Fig. 4A-C; purple point). In the landscape analogy it represents the lowest point in the ridge that separates the p3 and pMN valleys (Fig. 4A). Simulations of the SDE model indicated that intrinsic fluctuations around the pMN state are initially directed away from the transition point in WT. By contrast, in the Pax6 mutant fluctuations are oriented directly towards the transition point. As a consequence, fluctuations of the same magnitude would be more likely to reach the transition point in Pax6^−/−^ than WT cells. To characterise this rigorously, we calculated the most likely gene expression trajectory that a stochastic transition caused by fluctuations in gene expression will take between the pMN and p3 steady states. This path is obtained as the minimum of an “action functional” - the minimum action path (MAP, see Supplemental Section B). It provides a portrait of the dynamical landscape underlying a noise induced transition and is an analytical representation of the behaviour that can be observed in simulations (Fig. 4A & Supplemental Section C) [Perez-Carrasco et al., 2016, Kleinert, 2009, Bunin et al., 2012]. Consistent with the SDE simulations, in WT, the MAP from the pMN to p3 steady state does not follow the shortest route leading to the transition point. Instead, the levels of Pax6 drop rapidly and pitch away from the transition point, resulting in a curvature of the gene expression path between steady states (Fig. 4B). By contrast, in the absence of Pax6, the MAP is directly oriented towards the transition point (Fig. 4C). Taken together, the analysis suggests that the GRN affects the precision of a domain boundary by determining the dynamical landscape, without changing the level of noise in overall gene expression.

**Figure 4:**
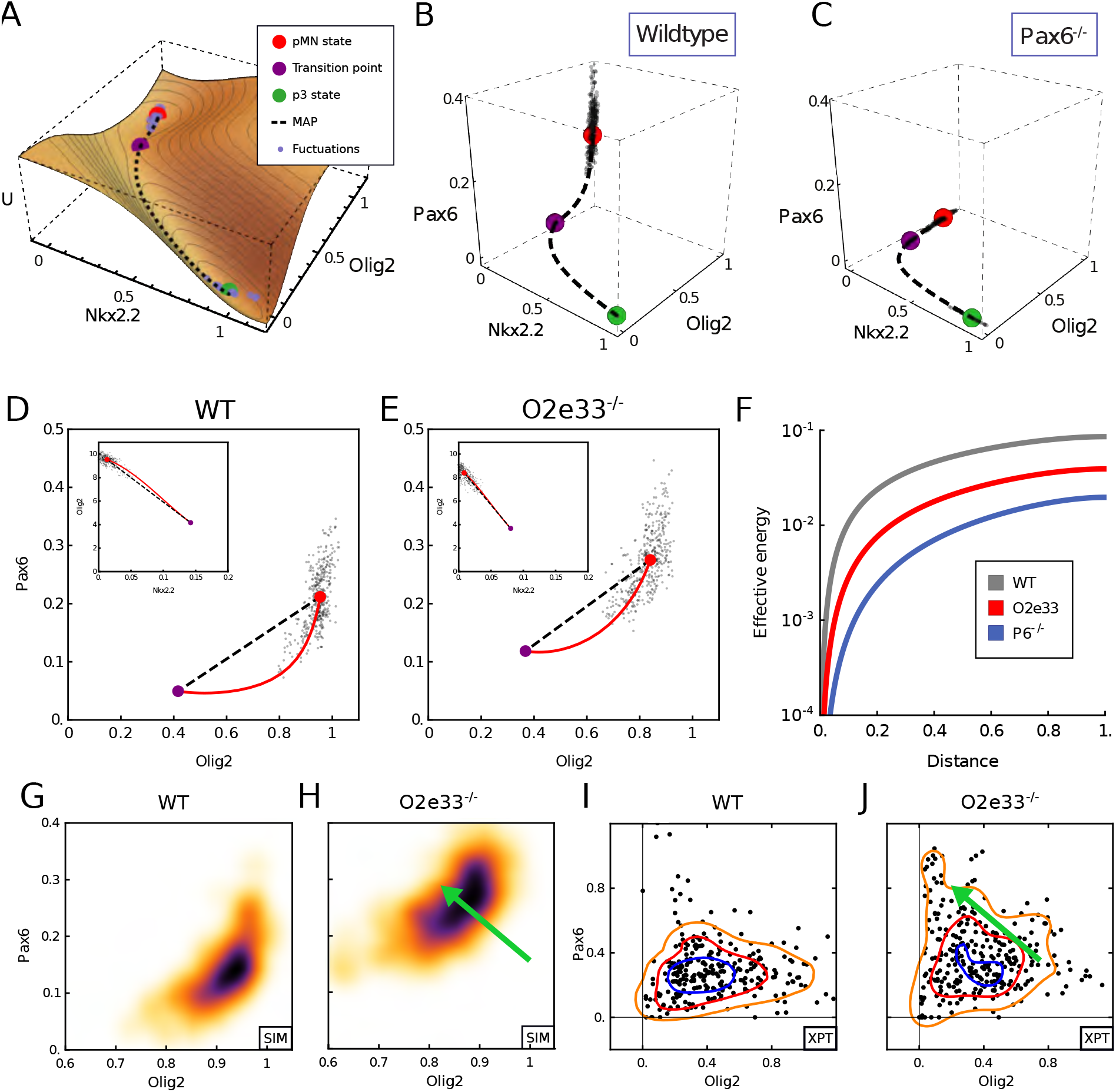
Mutant phenotypes affect the configuration of gene expression fluctuations. (**A**) A quasi-potential (U) representation of the neural tube dynamical system in region where noise driven transitions result in heterogeneity between pMN and p3 states. (**B-C**) Gene expression space view of the transition path from pMN (red point) to p3 (green point) steady states via the transition point (purple point). Simulated trajectory (dots) shows stochastic fluctuations from the pMN steady state. Axes show relative expression levels. WT (left) and Pax6^−/−^ (right) for neural tube position at fraction 0.1 of total neural tube length dorsal to the bifurcation point. (**D-E**) Projection into Olig2-Pax6 gene expression space of the minimum action path (red) predicted from the model and simulated trajectory (dots) in WT (I) and O2e33^−/−^ (J) at the same position as G-H. Insets show projection onto Nkx2.2-Olig2 axes. (**F**) Effective energy barrier (cumulative action) for noise-induced transitions, plotted along the transition path (normalised to unit length) at the same neural tube positions as G-J. WT (grey) has a higher barrier than O2e33^−/−^ (red), leading to longer jump times; O2e33^−/−^ in turn has a higher barrier than Pax6^−/−^ (blue). (**G**-**H**) Simulated Pax6 and Olig2 expression levels (black dots) for WT and O2e33^−/−^ in regions proximal to the p3-pMN boundary. (**I**-**J**) A shift to higher levels of Pax6 and reduced levels of Olig2 is observed in cells from O2e33^−/−^ mutants *in vivo* compared to controls. Axes show fluorescence intensity (arbitary units). Contour lines correspond to densities of the distribution of points, 0.6 (Orange), 1.6 (Red) and 2.6 (Blue).

For the O2e33^−/−^ mutants the MAP from pMN to p3 curved away from the shortest path to a lesser extent than for the WT; stochastic simulations further confirm this behaviour (Fig. 4D,E). Thus, in the absence of the O2e33 enhancer, stochastic fluctuations around the pMN steady state tended to take the system closer to the transition point than similar magnitude fluctuations in WT, making a noise driven switch in fate more likely in the mutant. Nevertheless, the curvature in the path in the O2e33^−/−^ system was greater than in the Pax6^−/−^ system, providing an explanation for the greater imprecision in Pax6^−/−^ embryos compared to the O2e33^−/−^ mutant (Fig. 4B-E).

To explore this phenomenom further, we calculated the action along the path for each genotype [de la Cruz et al., 2018](Fig. 4F & Supplemental Section C). This represents the effective energy required to reach a point along the transition path and is a measure of the extent of the barrier that has to be overcome for a fate transition. Consistent with the results of the simulations, the effective energy necessary for a noise induced transition was greatest for WT, less for O2e33^−/−^, and lowest for the Pax6^−/−^ mutant. Moreover, the analysis indicated that the initial part of the trajectory presented a more significant barrier to noise induced transitions in the WT than O2e33^−/−^ and Pax6^−/−^ mutants (Fig. S6A), corresponding to the relative divergences of their transition trajectories from the shortest route to the transition point.

An experimental testable signature of the alteration in the dynamical landscape in O2e33^−/−^ mutants would be changes in the relative expression levels of Olig2 and Pax6 in individual cells. In cells close to the pMN-p3 boundary O2e33^−/−^ mutants are predicted to have higher levels of Pax6 and lower levels of Olig2 than WT (Fig. 4G,H). We therefore compared single cell immunofluorescence in the boundary region of WT and O2e33^−/−^ embryos (Fig. 4I,J & Supplemental Section G.8). Consistent with the predictions, O2e33^−/−^ mutants had higher levels of Pax6 and lower levels of Olig2 than WT. Thus the experimental evidence supports the idea that the strength of regulatory interactions encoded in the GRN contributes to the precision of domain boundaries by configuring the dynamical landscape of the system to reduce the likelihood of a stochastic fluctuation resulting in a noise driven change in cell identity.

### A computational screen identifies mechanisms for precise boundaries

To ask whether other mechanisms could affect boundary precision, we performed a computational screen to identify three node networks capable of generating a sharp boundary in response to a graded input (Fig. 5A & Supplemental Section F). For the networks recovered from the screen, we compared the boundary precision with the extent the MAP deviates from the shortest path to the transition, a quantity that we refer to as “curvature” (Supplemental Section F). This showed a positive correlation, consistent with our observations in the WT network, of high curvature and low boundary width. This correlation supports the idea that the shape of the transition pathway contributes to boundary precision (Fig. 5C). Nevertheless, for any given level of boundary sharpness, there were a range of MAP curvature values. We therefore investigated additional features that might affect boundary precision. We found a subset of the networks do not rely on path curvature to achieve precision and instead functioned effectively as two node networks (Fig. 5D). For these networks, the major contributor to boundary precision was the rate at which the steady state and transition point separated in response to changes in level of the input signal: the higher the rate of separation, the sharper the boundary (Fig. 5B). We termed this “separation speed”. The most precise boundaries were generated by networks that exploited both separation speed and curvature, which includes the Pax6-Olig2-Nkx2.2 network (Fig. 5E-F).

**Figure 5:**
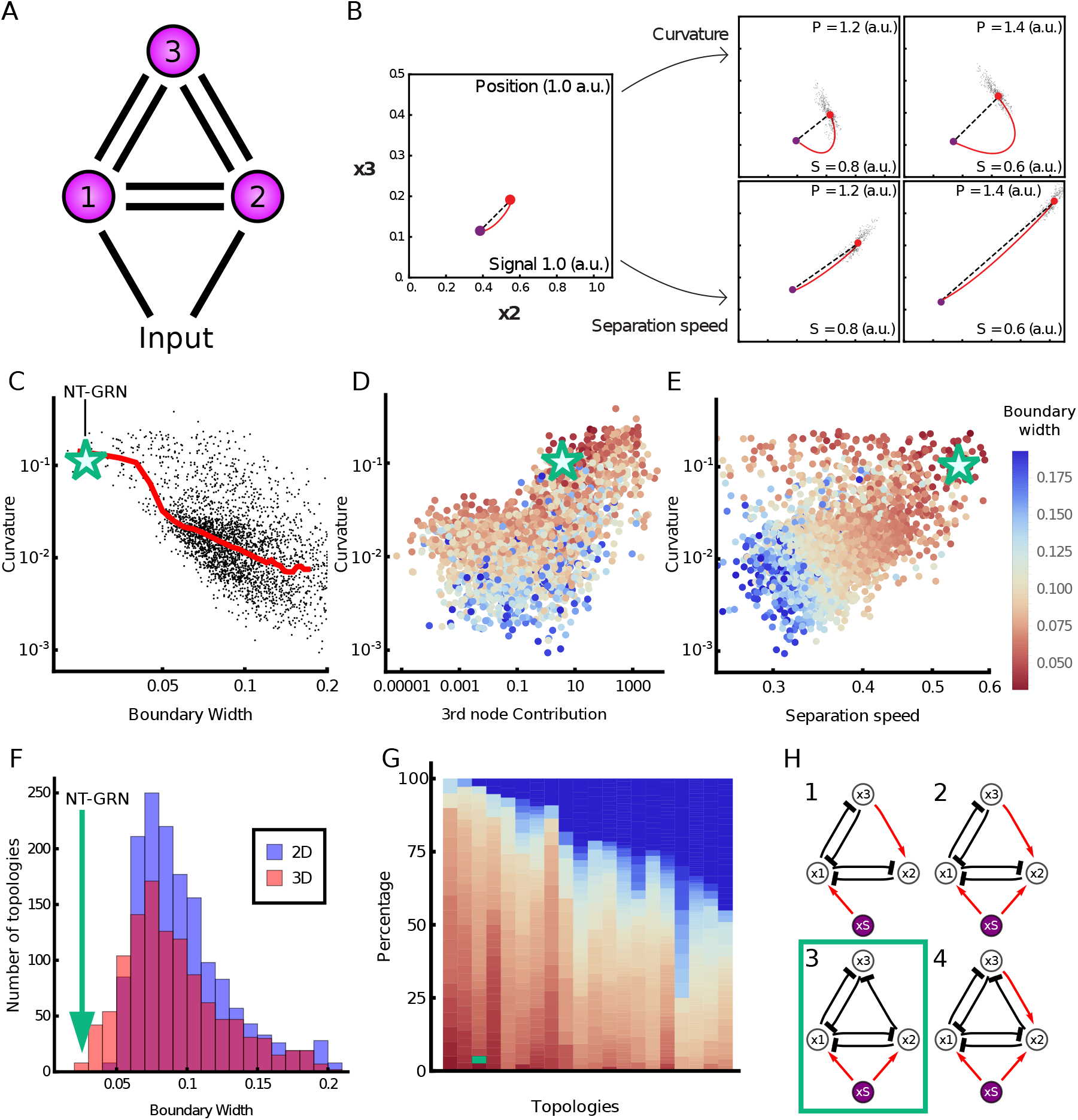
Computational screen reveals the design principles of precision. (**A**) Three node networks, comprising all possible interactions and a morphogen input into two nodes. (**B**) Two mechanisms for producing a precise boundary. Close to the boundary (Position 1.0 a.u.; Signal 1.0 a.u.) the steady state (red point) is near the transition point (purple point) in gene expression space. Further away (increasing Position; decreasing Signal) curvature of the MAP (red line) with respect to the shortest pathway (top row) or the rate at which the steady state separates from the transition point (bottom row) can contribute to increasing boundary precision. (**C**) For each network recovered from the screen (points), the boundary width was compared to the deviation of the MAP from the shortest path to the transition (curvature). Median value (red line) and illustrates that sharper boundaries (smaller width) tend to have higher MAP curvature. Green star represents the WT neural tube network. (**D**) Curvature compared to effective contribution of the third node in the network (boundary width indicated by colour of the point). (**E**) Curvature compared to separation speed. Colour of points by boundary width indicates both high curvature and high separation speed contribute to the sharpest boundaries. (**F**) Histogram of boundary width in 3D (red) and 2D (blue) networks. Green line represents the WT network. (**G**) The most common topologies, arranged in order of fraction of networks with precise boundaries; each column represents an individual topology. Dark blue indicates networks with a wider boundary. Topologies are shown in Fig. S16. (**H**) Four topologies that favour the sharpest boundaries. These networks comprise inhibition from node 2 to node 3, and lack repression from node 3 to node 2. The WT neural tube network has topology 3.

Finally, we assessed whether particular network topologies favoured boundary sharpness. Many topologies were able to generate sharp boundaries (Fig. 5G,H & Supplemental Section F), but four topologies appeared to be most effective (Fig. 5H). These tended to have similar separation speeds but much higher curvature than the networks with other topologies (Fig. S17). Crucial for this behaviour was the inhibition of *x*_3_ by *x*_2_ and the absence of repression of *x*_2_ by *x*_3_ (Fig. 5G & S16). This regulatory configuration generates curvature by allowing a steep decrease in *x*_3_, while sustaining high levels of *x*_2_ prior to the transition. Hence, an understanding of the dynamical properties of the GRN offers an explanation for its structure and the resulting gene expression behaviour that determines tissue patterning.

## Discussion

In this study we provide evidence that the spatial heterogeneity that results from the stochasticity of gene expression can be attenuated by the dynamics of the GRN to enhance the precision of gene expression in developing tissues. This mechanism does not rely on suppressing stochastic fluctuations in individual genes, nor on cell-to-cell communication, but instead configures the dynamical landscape of the regulatory network to increase the fidelity of decision making. This strategy - “precision by design” - highlights the capacity of gene regulatory circuits to contribute to robust tissue patterning and identifies a mechanism that might be exploited in other biological settings requiring precise responses from groups of cells.

### GRN dynamics contribute to precise boundaries without attenuating gene expression noise

Molecular noise is a universal feature of gene expression [Raj and van Oudenaarden, 2008, Raser and O’Shea, 2005, Chalancon et al., 2012]. Despite this, patterns of gene expression in developing tissues are remarkably reproducible and precise, as exemplified by the sharp boundaries of gene expression that delimit distinct domains of cells in many tissues. This spatial precision is critical for the accurate assembly of tissues. For example, along the anterior-posterior axis of the Drosophila embryo the expression of genes that partition the blastoderm into the major elements of the body axis are positioned with an accuracy of 1% or better [Dubuis et al., 2013, Petkova et al., 2019]. Similarly, in the central nervous system the correct positioning of different neuronal subtypes is a major determinant of their subsequent patterns of connectivity and underpins the formation of functional neural circuits [Jessell et al., 2011, Balaskas et al., 2019].

Mechanisms involving cell-cell interactions to correct initial imprecisions in the spatial organisation of tissues have received considerable attention [Xu et al., 1999, Standley et al., 2001, Rudolf et al., 2015, Dahmann et al., 2011, Addison et al., 2018]. Differential cell adhesion between neural progenitors with different cellular identities has been proposed to refine initially disordered patterns [Lei et al., 2004, Xiong et al., 2013, Tsai et al., 2020]. However, neither differential adhesion nor cell sorting appear to be the sole explanation for the precision of patterning in the neural tube. Lineage tracing in the mouse and chick neural tube [Kicheva et al., 2014, Leber and Sanes, 1995] indicates that sister cells form contiguous clones and there is no evidence that clones at a domain boundary behave in a way compatible with differential interactions across a boundary. Moreover, in both Pax6^−/−^ and O2e33^−/−^ mutants neural progenitors with distinct identities, pMN and p3, producing MNs and V3 neurons respectively, continue to be generated, but these different progenitor types intermix to a greater extent than normal. If the differential expression of cell adhesion molecules or different mechanical properties explained the sharpness of the boundary between pMN and p3 cells, this would lead to the sorting of pMN and p3 cells in the mutant embryos. Nevertheless, cell adhesion might play a role in the neural tube of teleosts [Xiong et al., 2013, Tsai et al., 2020]. Unlike the epithelial neural tube of amniotes, the zebrafish neural tube initially comprises unpolarised non-epithelial cells and sister cells disperse widely, including contralaterally, in the neural tube. This raises the possibility that differential cell adhesion plays a more important role in the anamniote neural tube.

Similar to many developing tissues, the neural tube is patterned by graded signals that are transformed into discrete cell identities by the downstream GRN acting as a series of toggle switches to produce discontinuous changes in cell identity across the tissue [Sagner and Briscoe, 2019]. Previous studies have explored how properties of extracellular patterning signals [Bollenbach et al., 2008, Tkačik et al., 2015, Lucas et al., 2018, Sokolowski et al., 2012, Zagorski et al., 2017] and features of the regulation of individual genes [Perry et al., 2010, Frankel et al., 2010, Lagha et al., 2012, Little et al., 2013, Battich et al., 2015, Dickel et al., 2018, Osterwalder et al., 2018, Paliou et al., 2019] can contribute to the fidelity of gene expression. Some of these mechanisms may play a part in the precision of neural tube gene expression. For example, paralogs of several of the key transcription factors are coexpressed in the neural tube and appear to function, at least partially, redundantly [Vallstedt et al., 2001, Holz et al., 2010]. In addition, the provision of antiparallel signaling gradients emanating from the opposing dorsal and ventral poles of the neural tube have been implicated in increasing the precision of gene expression in central regions of the spinal cord [Zagorski et al., 2017]. However the ventral regions of the neural tube where the p3-pMN boundary is positioned is out of range of the dorsal signal. The changes in boundary precision in the neural tube of the Pax6^−/−^ and O2e33^−/−^ mutants are not explained by changes in noise amplitude in individual genes or global changes in the magnitude of the noise. Instead the genetic perturbations we analysed alter the dynamics of the GRN and these change the configuration of gene expression fluctations and make noise driven transitions between cell states more likely. Thus the dynamics of the gene regulatory network affect patterning precision, without altering the stochasticity of individual components of the system, indicating that the configuration of gene expression noise, not simply the magnitude, affects development precision.

Stochastic fluctuations in gene expression are expected to result in variations in the position at which cells switch identity and produce indistinct boundaries. There is a trade-off between the steepness, precision and speed of boundary formation [Chalancon et al., 2012, Lv et al., 2014, Perez-Carrasco et al., 2016, Tran et al., 2018]. If gene expression were deterministic, a graded signal controlling such a switch would generate a sharp, precisely positioned gene expression boundary in the tissue. However, the effect of stochastic fluctuations is that an increase in non-linearity and switch-like behaviour decreases boundary precision: stochastic fluctuations generate a change in gene expression that is independent of changes in signal input. Our analysis of the Pax6-Olig2-Nkx2.2 network revealed that the GRN is configured to decrease the probability of such spontaneous noise driven transitions while retaining the ability to produce discontinuous switch-like changes in gene expression, thereby generating a sharp, precise boundary in the tissue. This mechanism enhances boundary precision even in the presence of noise in the signalling gradient (Supp. F & Fig. S18). Moreover, the same regulatory mechanism that decreases the probability of a noise driven transition from pMN to p3 also produces hysteresis, this ratchet-like effect means that once a cell has adopoted a p3 identity it is unlikely to transition back [Balaskas et al., 2012]. Thus the dynamics of this GRN increase the precision of the pMN-p3 boundary by decreasing the probability of transitions between pMN and p3 in either direction.

### Configuring the dynamical landscape to maximise precision

. Viewed from the perspective of the Waddington landscape [Waddington, 1957], spontaneous changes in cell state resulting from gene expression fluctuations would be represented as a cell being displaced from one valley to another by traversing the intervening ridge (Supplementary Section B). The dynamical landscape produced by the Pax6-Olig2-Nkx2.2 network is configured so that the height of the ridge between the two valleys changes rapidly as the level of morphogen signalling changes. This is evident from analysis of the MAP, which reveals that transition trajectories between cell states diverge substantially from the shortest route to the transition point (Fig. 4B-E). The consequence of this is that the effective energy necessary for a noise induced fate transition was higher for WT than either of the mutants with a perturbed GRN (Fig. 4F & Supplemental Section C). Thus the GRN minimizes the range of signalling for which noise induced transitions are likely to occur, without altering the stochasticity of individual genes, hence increasing boundary sharpness.

This mechanism, which we termed “curvature”, was identified in an unbiased computational screen of three node networks responding to a graded input signal (Fig. 5). In addition, the screen recovered a second mechanism - “separation speed” - that relied on the rate at which the two cell states separated in response to changes in the level of input signal (Fig. 5B). In the context of the Waddington landscape, separation speed can be viewed as changes in signal levels producing rapid changes in the distance between the two valleys. A feature of this second mechanism is that it can be implemented with only two genes. However, instead of producing two cell states both with uniform levels of gene expression, one of the resulting cell states is characterised by a gradient of gene expression S15). This might limit its utility in some tissue patterning roles. By contrast, the curvature mechanism requires a minimum of three nodes to implement, but it is able to produce two cell states with almost constant levels of gene expression. Nevertheless, the two mechanisms of speed and curvature are not mutually exclusive and the networks recovered by the screen that generated the most precise boundaries combined both mechanisms.

### Regulatory principles of patterning precision

Similar to other recent studies [Cotterell and Sharpe, 2010, Schaerli et al., 2014, Verd et al., 2019], the screen indicated that the dynamics of the networks, not simply the network topology, were key to determining the resulting precision. A feature shared by many of the networks with the sharpest boundaries, including the neural tube network, was an asymmetry in inhibition between two of the genes (Fig 5H). Specifically, *x*_3_ (Pax6) repressed *x*_2_ (Olig2), but not vice versa. Moreover, the graded expression of Pax6 (*x*_3_) within the domain is indicative of the separation speed mechanism, providing evidence that this too contributes to boundary precision while allowing uniform levels of Olig2 expression (*x*_2_; the gene necessary for defining the identity of this domain). This analysis therefore raises the possibility that the dynamics of the Pax6-Olig2-Nkx2.2 network were adopted in the developing vertebrate neural tube for its capacity to generate distinct cell type identities with precise boundaries. In this context, it is striking that gene circuits with similar structure and dynamics have been implicated in the patterning of the anterior-posterior axis of the Drosophila embryo [Akam, 1987, Ingham, 1988, Sánchez and Thieffry, 2001, Manu et al., 2009, Verd et al., 2017] and the Drosophila eye imaginal discs [O’Neill et al., 1994, Rebay and Rubin, 1995, Graham et al., 2010] (Supplemental Section F & Fig. S19, S20). Taken together therefore, the computational screen defines design features of multi-stable gene circuits that are suited to the generation of sharp boundaries in response to graded inputs.

The dynamics of a GRN are governed by the strength of regulatory interactions between the components of the network, which in turn are determined by cis regulatory elements and their binding to transcription factors [Davidson, 2010]. Many genes involved in development are associated with two or more cis regulatory elements that seem to function in a partially redundant manner [Perry et al., 2011, El-Sherif and Levine, 2016, Cannavò et al., 2016, Dunipace et al., 2019]. This appears to be the case for Olig2, where removal of the O2e33 element perturbs, but does not completely abrogate, Olig2 expression (Fig. 2). This supports the idea that one function of multiple cis regulatory elements is to provide robustness and precision to gene expression [Perry et al., 2010, Frankel et al., 2010, Lagha et al., 2012, Battich et al., 2015, Dickel et al., 2018, Osterwalder et al., 2018, Paliou et al., 2019, Tsai et al., 2019]. Our analysis indicates that these functions are not simply a consequence of multiple elements supplying duplicate activities. Instead individual cis regulatory elements provide specific dynamical properties to gene regulation that sculpt the gene expression landscape. Thus, distinct cis regulatory elements of a target gene serve specific dynamical functions within a GRN.

Taken together, our analysis illustrates how a tissue level feature - the spatial precision of gene expression patterns - is influenced by cell autonomous mechanisms, implemented by cis regulatory elements that influence the activity of a network of interacting transcription factors. The data reveal that the potential detrimental effects of stochastic fluctuations in gene expression that would lead to spatial heterogeneity can be attenuated by the dynamics of the GRN. We term the strategy “precision by design” as it arises from the integrated function of the gene circuit and is not intrinsic to any individual network component. This provides insight into decision making in multicellular systems and highlights how an understanding of the dynamics of GRNs can explain its structure and function. More generally, identifying the principles that produce robust and precise outputs despite the inherent stochasticity of gene expression should assist in the future design, modification and engineering of gene circuits.

## Acknowledgements

We thank JP Vincent, B Verd and members of the lab for constructive discussions and comments. We are grateful to the Flow Cytometry, Biological Resource and HPC Facilities of the Francis Crick Institute. This work was supported by: the Francis Crick Institute, which receives its core funding from Cancer Research UK (FC001051), the UK Medical Research Council (FC001051), and Wellcome (FC001051); funding from Wellcome [WT098325MA and WT098326MA]; the European Research Council under European Union (EU) Horizon 2020 research and innovation program grant 742138. RPC acknowledges the UCL Mathematics Clifford Fellowship.

## Author Contributions

KE, EHD, PS & JB conceived the project, interpreted the data and wrote the manuscript with input from all authors. KE performed all experiments except those listed under other authors. EHD performed theoretical modelling and data analysis. LGP performed the protein copy number quantifications. RPC contributed to building the mathematical models and provided advice. AS generated the Olig2-T2A-mKate2 ES cell derived neural progenitors. VM analysed the ATAC-seq data and supervised experimental work. PS & JB supervised the project.

## Supplementary Material

## A Supplementary Figures

**Figure S1:**
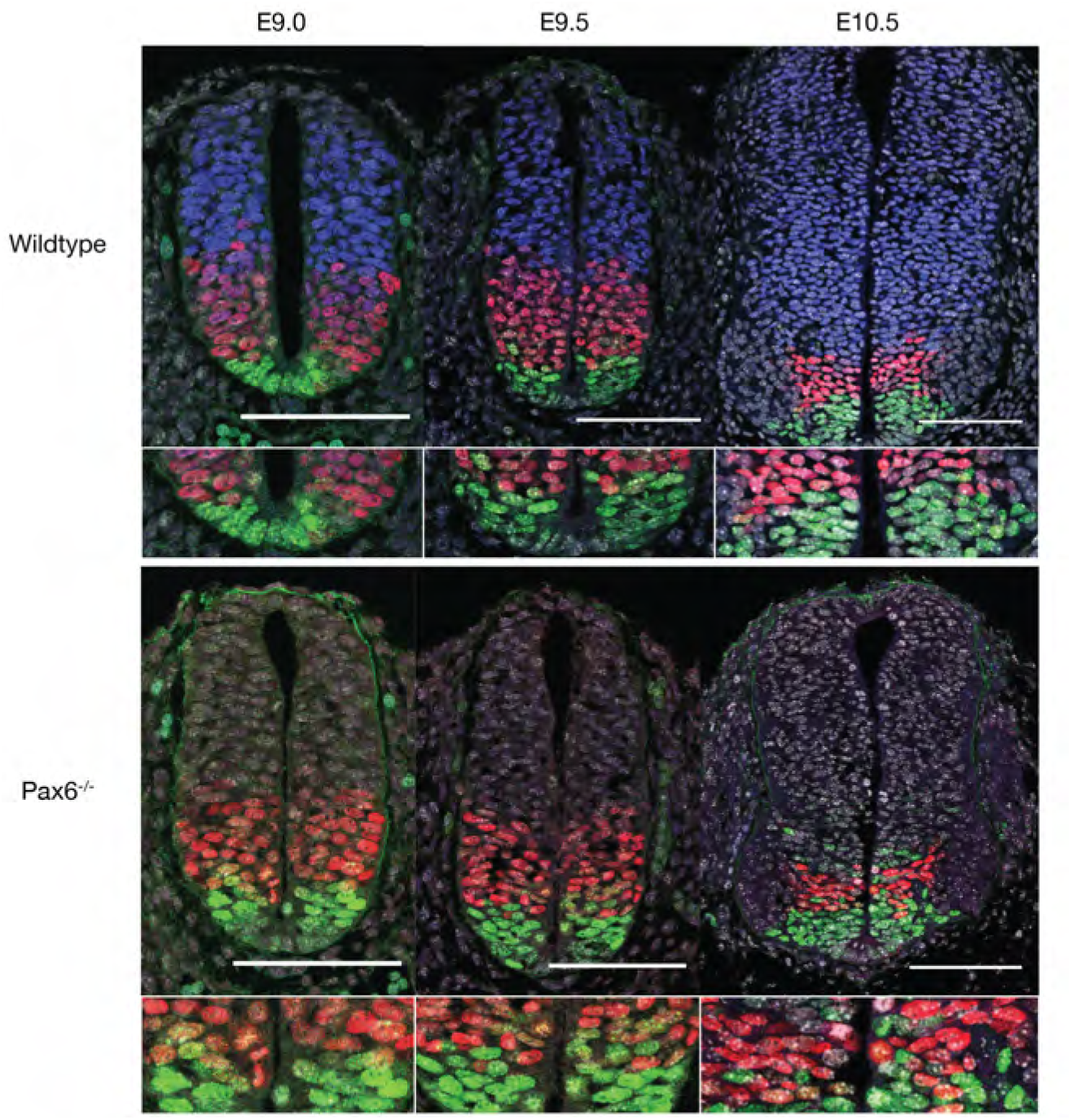
pMN-p3 boundary precision decreases over time in Pax6 mutants. Transverse sections of wildtype and Pax6^−/−^ embryos between e9.0 and e10.5 stained for Pax6 (blue), Olig2 (red) and Nkx2.2 (green). Scale bar = 100*μ*m. The pMN-p3 boundary becomes less well defined at later time points.

**Figure S2:**
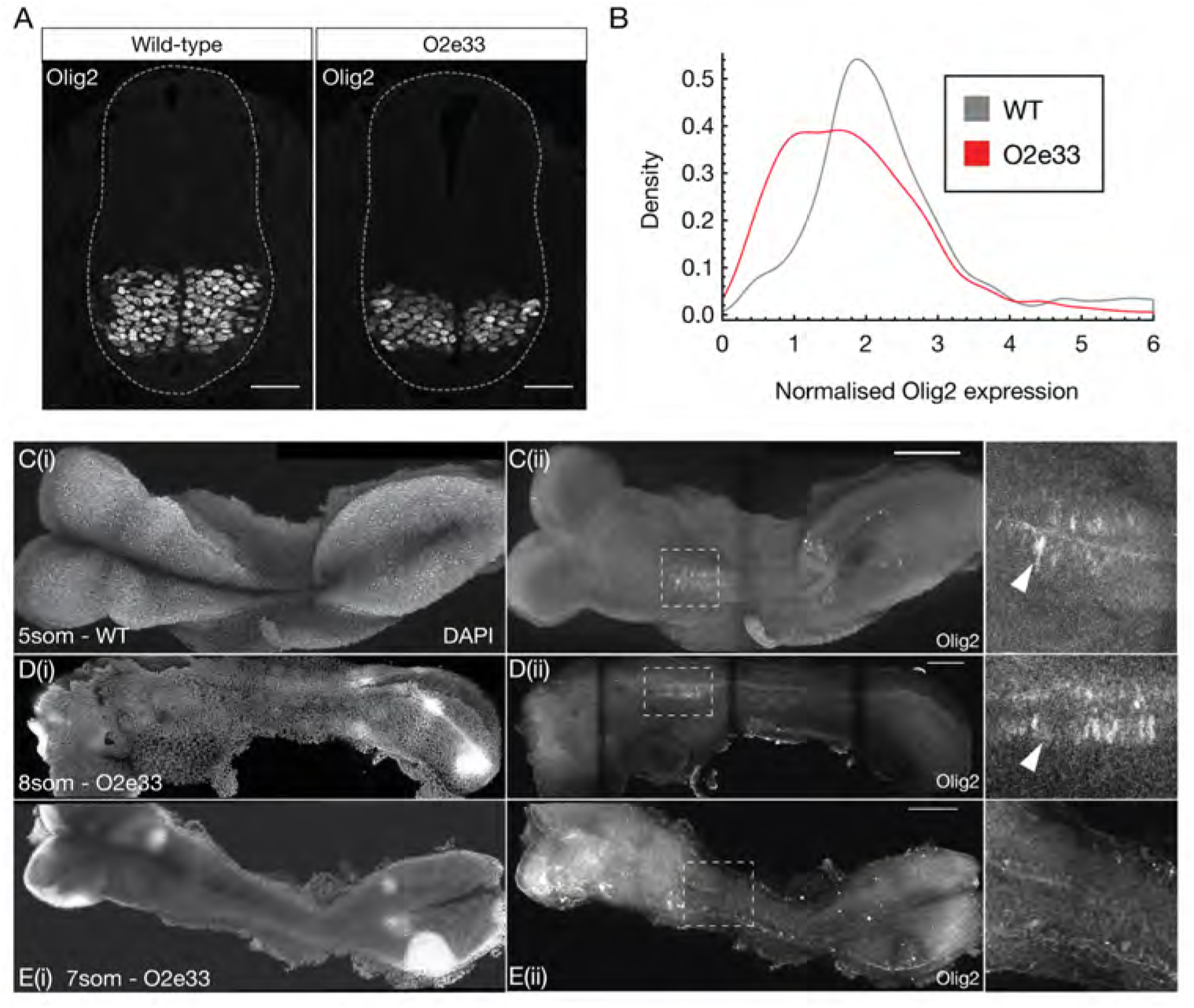
Olig2 expression in O2e33 ^−/−^ mutants is lower and delayed in onset. (**A**) Transverse brachial sections of e9.5 WT and O2e33^−/−^ embryos stained for Olig2. The O2e33^−/−^ embryo has a smaller Olig2 domain with reduced expression levels. Scale bar = 50*μ*m (**B**) Normalised Olig2 expression for single cells in WT and O2e33^−/−^ embryo sections. (**C, D, E**) Wholemount images of WT (C) and O2e33^−/−^ mutants (D, E) for DAPI (i) and Olig2 staining (ii-iii). Expression of Olig2 in wildtype is observed at 5 somites but in O2e33^−/−^ Olig2 onset occurs later at 8 somites. Olig2 is not observed in O2e33^−/−^ embryos at 7 somites. Scale bar = 100*μ*m

**Figure S3:**
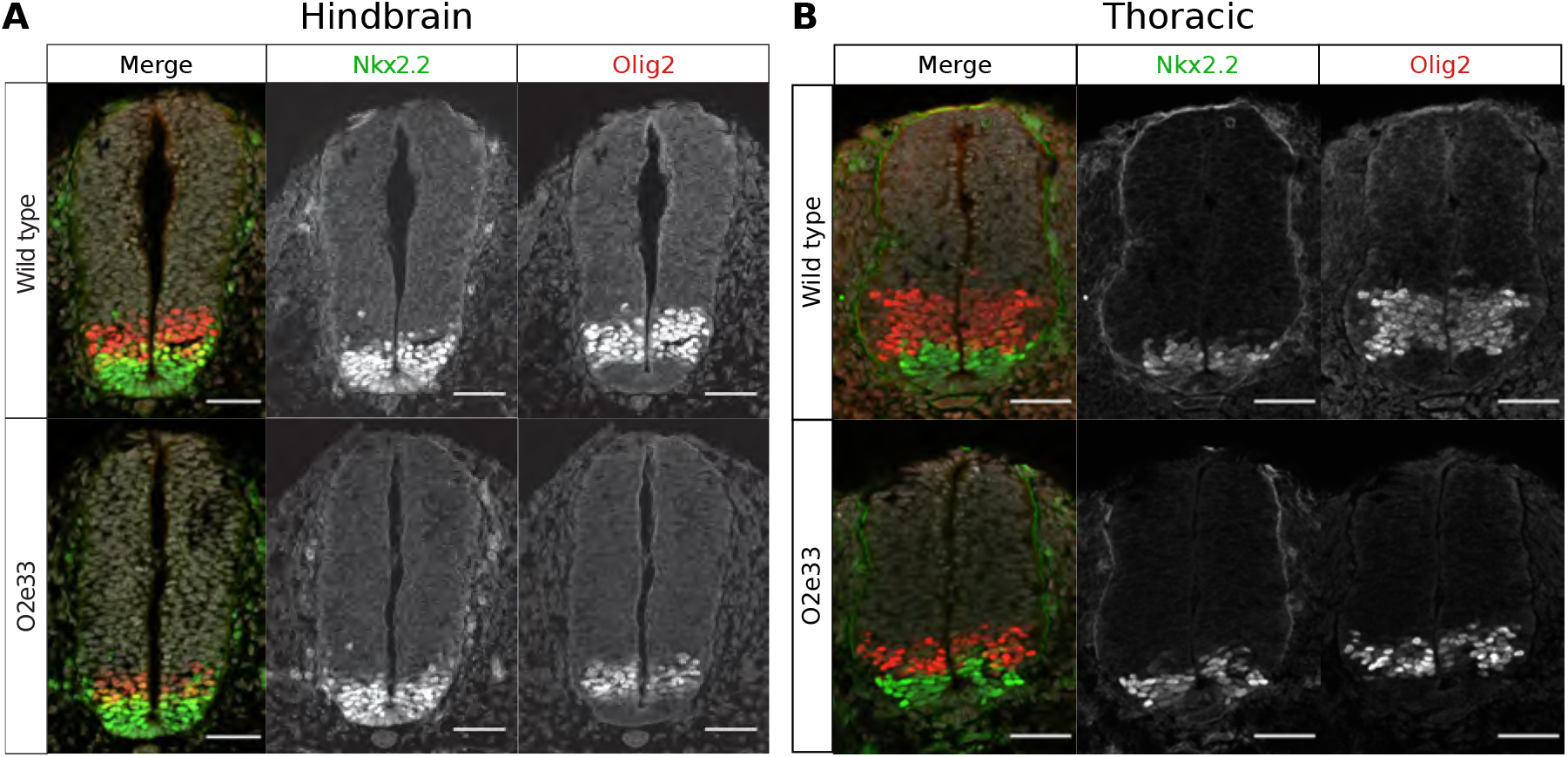
Transverse sections of the hindbrain (A) and thoracic region (B) e9.5 wildtype and O2e33^−/−^ embryos. Stained for Olig2 (red) and Nkx2.2 (green). Scale bar = 50. (A) Hindbrain: The pMN domain is smaller and the pMN-p3 boundary is less well defined in O2e33^−/−^ mutant embryos. (B) Thoracic region: The pMN domain is smaller and there is more intermixing between pMN and p3 cells in O2e33^−/−^ mutant embryos.

## B Glossary of dynamical systems terminology

The terms in this glossary come from the field of dynamical systems theory and more detail can be found in [Kuznetsov, 2008, Strogatz, 2014]. We then also use elements relating to stochastic processes for which further information can be found in e.g. [Van Kampen, 2007].

- **Deterministic system** Deterministic systems are those that involve no randomness and will therefore *always* behave in the exact same way when started from the same conditions. In this study deterministic systems model the production and degradation of genes in the absence of any stochasticity.
- **Stochastic systems** These are systems of equations that incorporate randomness such that the system *will not* behave the same way every time. In our study, these are derived from the **deterministic system** by adding a stochastic element to form a **Chemical Langevin Equation**.
- **Chemical Langevin Equation** The Langevin equation was derived by Paul Langevin as an equation that approximates the randomness generated by individual processes and has been adapted to describe chemical reaction systems [Lemons and Gythiel, 1997, Gillespie, 2000]. It assumes each individual reaction in a system takes place with Gaussian noise and has been shown to be accurate for systems in which the number of molecules for each component (*e.g.* transcription factor) in the system is sufficiently large. It involves incorporating **stochastic terms**, which describe the noise, into the **deterministic system**.
- **Phase space** Phase space is an abstract space in which each dimension represents the concentration of one of the components (transcription factors) of the gene regulatory network. This allows the dynamics of the GRN to be visualised geometrically, such that the change in concentration of the TFs over time traces out a line in phase space.
- **Critical points** A critical point is a point in **phase space** where the **deterministic system** does not change over time. That is, the time derivative of all concentrations at a critical point is zero. These points can represent **stable fixed points** of a system, or **unstable fixed points**.
- **Stable fixed points** (Attractor points) A stable fixed point is a type of **critical point**. If in the immediate surroundings of a critical point the dynamics of the **deterministic system** indicate that the system moves towards the fixed point from any direction, this critical point is termed a stable fixed point. This notion is referred to as Lyapunov stability [Lyapunov, 1992]. In a “Waddington-like” landscape visualisation, stable fixed points can be thought of as the basins at the bottom of valleys. In this study, we look at systems with a maximum of two **stable fixed points** (Fig. S4 & S5).
- **Unstable fixed points** An unstable fixed point is also a type of **critical point**. In contrast to **stable fixed points**, if the analysis of the **deterministic system** shows that the system moves away from the fixed point when started some small distance away in at least one direction, the point is termed an unstable fixed point. This means that the system will only remain at this point if it is located there *exactly*. A **stochastic system** will not remain at such a point as the stochastic terms will eventually result in the system moving away along an unstable direction. In a “Waddington-like” landscape, unstable fixed points can be thought of as peaks or ridges from which the cell will move away.
- **Saddle point** (Transition point) A saddle point is a type of **unstable fixed point** that is attractive in at least one dimension. In the systems within this study (as in many others), saddle points separate **stable fixed points**. A system will approach a saddle point and pass through it during a transition between **stable fixed points**. In a “Waddington-like” landscape, a saddle point appears like a mountain pass between two peaks, or a saddle, hence the name (Fig. S4 & S5).
- **Bifurcation point** In the systems in this study bifurcation points are the positions along the morphogen gradient where the system goes from having a single **stable steady state** to being bistable; this means having two **stable steady states** and one **saddle point** (Fig. S4).
- **Fluctuations in concentration** In a **stochastic system** the concentrations of the molecules fluctuate *at all times* in a way described by the **Chemical Langevin Equation**. This means that a system never stabilises at a constant concentration, even at a **stable fixed point**. Fluctuations in concentration around a stable fixed point remain and can be analysed and visualised in **phase space** (Fig. S5).
- **Noise driven transitions** **Fluctuations in concentration** near a **stable fixed point** move the state of the system away from the **stable fixed point** in phase space. This can result in the system reaching a **saddle point** and as a consequence transitioning from the original **stable fixed point** to the basin of attraction of a different **stable fixed point**. This process is a noise driven transition (Fig. S4 & S5).
- **Minimum Action Path (MAP)** From the equations for the **stochastic system** it is possible to calculate the most likely path that a system will take to complete a transition from one **stable fixed point** to another [Kleinert, 2009, Bunin et al., 2012]. This is termed the Minimum Action Path (MAP) and can be visualised as a gene expression trajectory in the **phase space** of TF concentrations. In a “Waddington-like” landscape visualisation such paths can be thought of as the lowest paths in the landscape, which cells are most likely to follow as they move from one state to another(Fig. S5).

In addition, we use the following terms to describe the characteristics of a dynamical system that contribute to precise boundaries.

- **Curvature** This is a measurement of how directly the **Minimum Action Path (MAP)** connects a **stable fixed point** to a **saddle point**. In **phase space**, the length of the **MAP** is compared with the shortest distance (straight line) between the initial **stable steady state** and the **saddle point**, at a fixed neural tube position. The greater the ratio between these two distances, the higher the **curvature**.
- **Separation speed** A measurement of the Euclidean distance in **phase space** between the **stable fixed point** at which the system starts and the **saddle point**, at a fixed neural tube distance from the **bifurcation point** (the same neural tube position where also the **curvature** is determined). This is termed the separation speed as it indicates how fast the **stable fixed point** and **saddle point** separate in response to distance from the source of morphogen.

**Figure S4:**
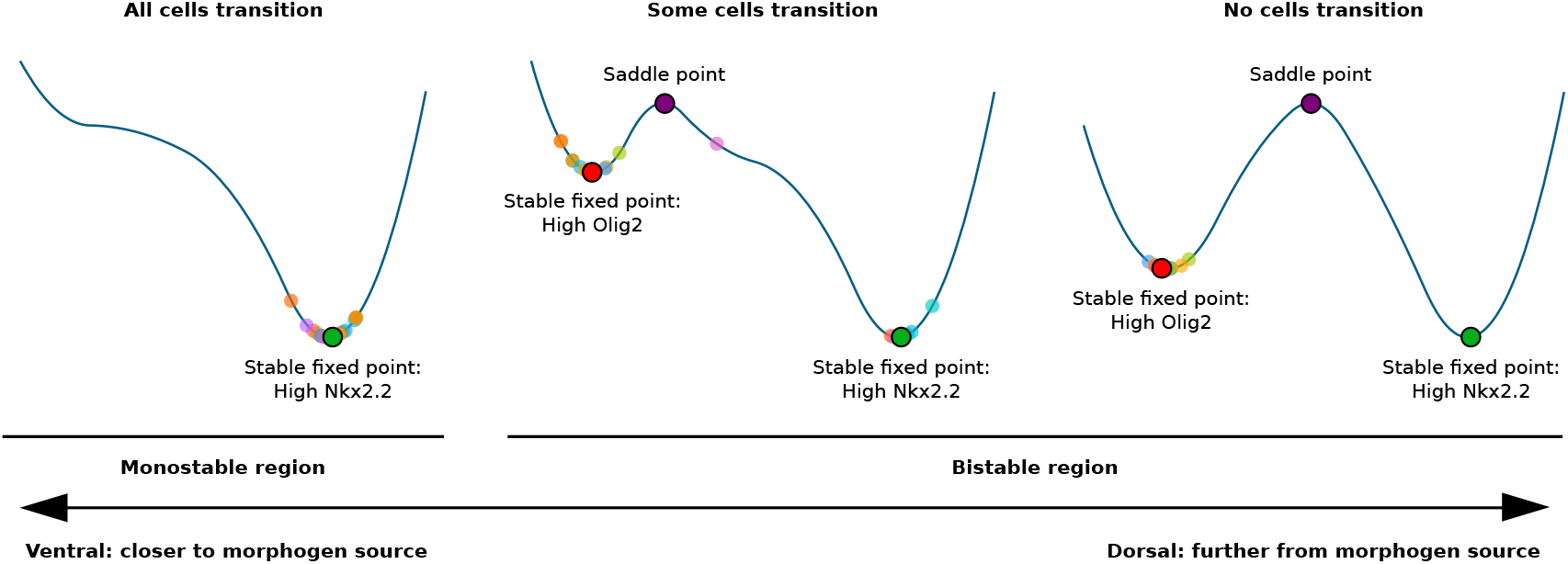
One-dimensional sketches of the dynamical landscape of the neural tube network at multiple dorso-ventral positions as indicated by the bottom arrow. The larger dots with black contours indicate **critical points** as labelled on the plot. The multiple smaller dots represent the final gene expression profile of different simulations of **stochastic** GRN at the same neural tube position. The stochastic nature of the systems leads to cells not following the same identical path. The leftmost plot indicates monostability for high Nkx2.2 near the ventral end of the neural tube; here all cells present high levels of Nkx2.2 as there is no other stable fixed point. In the bistable region all systems start out with high levels of Olig2 as happens in the neural tube. The middle plot represents the landscape slightly dorsal to the **bifurcation point**. The system here presents bistability so that **noise driven transitions** can occur where the system is driven to and beyond the **saddle point**; however these transitions *do not always occur*, leading to heterogeneity in fate decisions for cells at this position. The plot to the right represents cells much further dorsal of the bifurcation; here the probability of a system reaching the **saddle point** is extremely low even with stochastic terms as can be appreciated from the figure. In this region despite the existence of bistability, only the high Olig2 **stable fixed point** is observed as the probability for **noise driven transitions** to occur is vanishingly small.

**Figure S5:**
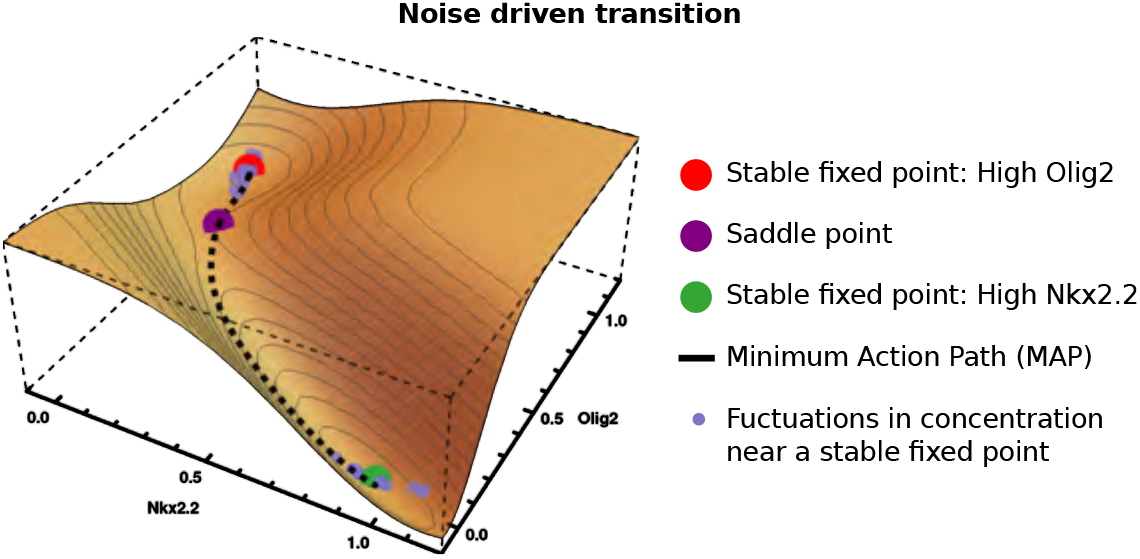
Representation of the dynamical landscape of the neural tube network at a region of bistability, slightly dorsal to the **bifurcation point**. The *x* and *y* axes are the concentrations of Nkx2.2 and Olig2 respectively (2D phase space), whereas the *z* axis represents the landscape of the system. The plot relates to the same neural tube position as the middle figure in Fig. S4, therefore there is heterogeneity in fate decisions. The colouring of the **critical points** is consistent with Fig. S4; see also the legend. The light blue dots represent two different simulations near each **stable fixed point**, illustrating **fluctuations in concentration**. The thick black line illustrates the **MAP** from the high Olig2 **stable fixed point** to the high Nkx2.2 **stable fixed point**. Note that it passes through a critical point – a saddle (purple dot).

## C Formulation and analysis of stochastic GRN dynamics

### Formulation of stochastic dynamics

In order to investigate heterogeneity of gene expression in the neural tube we made use of stochastic differential equations that describe the GRN and in particular the time evolution of the concentration *x*_*j*_ of each TF *j*. We start with a thermodynamic-like model as detailed in [Cohen et al., 2014], which captures the macroscopic behaviour by a system of ODEs; these contain terms for production and decay of each TF. The ODE description corresponds to the limit of a reaction volume Ω that is large enough for the copy numbers Ω*x*_*j*_ of all protein species to be large, allowing fluctuations to be neglected; formally one takes Ω → ∞. When Ω is finite, stochastic effects occur. These can be described by the chemical Langevin equation, a system of SDEs, see e.g. [Van Kampen, 2007, Gillespie, 2000]. The drift, *i.e.* the systematic variation with time in the SDEs coincides directly with the deterministic limit. The diffusion (stochastic) term arises from the stochastic nature of the individual protein production and decay reactions; it is a Gaussian white noise [Gillespie, 2000] whose covariance structure is determined by the mean reaction rates. In our case the chemical Langevin equation for the protein levels *x*_*j*_ within the GRN takes the form:

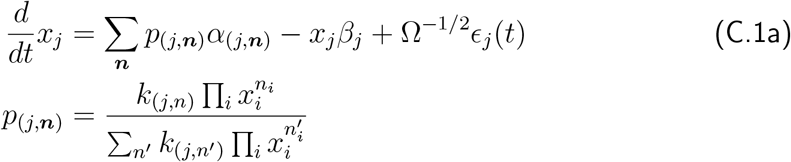

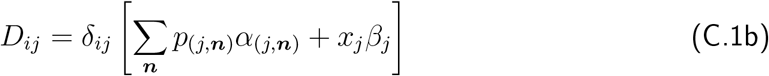

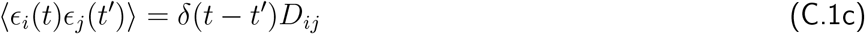

The deterministic part of these equations is equivalent to those used in [Cohen et al., 2014]. The covariance (C.1b,C.1c) of the zero mean Gaussian white noise *ϵ*_*j*_(*t*) arises from the decay and production of each protein being independent and random, given the concentration of the regulators of the relevant gene. In the equations above, *α* represents protein production rate and *β* degradation rate, while the *w* provide the weights of the respective DNA conformations (*j*, ***n***) when multiplied by the respective concentration. The conformations are labelled by the protein *j* being produced and the numbers ***n*** = {*n*_*i*_} of TF molecules bound. The *δ* in (C.1b) and (C.1c) are the Kronecker and Dirac delta respectively. As explained above, Ω is the volume of the system in which all reactions take place.

When looking at the chemical Langevin equation (C.1a), one notices that the rate ∑_***n***_*p*_(*j*,***n***)_*α*_(*j*,***n***)_ for producing protein *j*, has a nonlinear dependence on the TF concentrations *x*_*i*_. One might be concerned that with such a nonlinear dependence, modelling production of protein *j* as a single reaction is too simplistic. However, (C.1a) can be obtained from a larger system of simple unary and binary mass action reactions, in which the concentration of each DNA conformation is kept track of individually. We only sketch this construction here and explain its implications for the stochastic terms in (C.1a); for further details see [Herrera-Delgado et al., 2018]. The deterministic part of the time evolution of the DNA concentrations is given as follows:

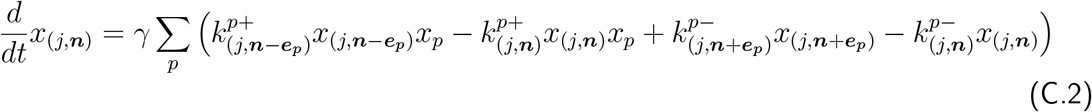

Here 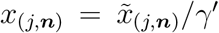 tracks the concentration of each DNA conformation and is scaled down by a large factor *γ*′ to account for the low quantity of binding sites in relation to protein numbers. Correspondingly the protein production rate constants 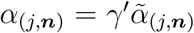 have to be large in order to give an appreciable overall rate of protein production nonetheless.

To derive the correct stochastic equations for the protein species, the large *γ*-limit of (C.2) is taken: the concentration of each DNA conformation then changes sufficiently quickly that it constantly tracks the instantaneous protein concentrations. For appropriately chosen binding and unbinding rate constants 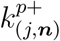 and 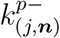 this leads back to the thermodynamic-like form of the *deterministic* part of the protein dynamics in (C.1a) [Herrera-Delgado et al., 2018]. As shown in [Thomas et al., 2012] the existence of fast species (in our case, DNA conformations) can lead to additional terms arising in the *noise* acting on the slow species (protein production), as a consequence of reactions between slow and fast species. In our case it turns out that these extra noise terms scale with *γ*′/*γ*. We then make use of the biological meaning of the terms: 1/*γ* represents the timescale of reaction rates for TF binding to DNA and 1/*γ*′ represents the characteristic time for the process of going from active DNA to producing a protein. We find it biologically reasonable to choose a 1/*γ* that is substantially smaller than 1/*γ*′, given the many biological processes necessary for the production of a fully functional protein. The ratio *γ*′/*γ*is then small so that the additional noise terms that arise from the general calculation in [Thomas et al., 2012] become negligible, leaving exactly the noise terms in (C.1c). The intuition is that because protein production is slow compared to binding and unbinding of factors to DNA, noise from the many binding and unbinding events during production averages out; the overall noise then arises only from the stochasticity of the production processes, at the relevant average DNA concentrations. We note that in accordance wSit1h1this conclusion, explicit calculations show that when *γ*′ is of the order of *γ* or larger, additional noise terms from the stochasticity in DNA concentrations do enter the dynamics of the protein concentrations. Moreover, these additional terms are dependent on the precise choices of binding and unbinding rates, which are only partially constrained by the requirement that the thermodynamic-like deterministic equations (C.1a) are retrieved for large *γ* [Herrera-Delgado et al., 2018].

### Amount of noise

The noise level in our model is set by Ω^−1^, the inverse reaction volume. This determines the scale of the stochastic fluctuations in protein production and decay, both of which the model represents as single step processes. A larger Ω thus leads to smaller stochastic effects. In equation (C.1a), multiplying Ω by the concentration of a protein species gives the number of molecules for that protein. In our calculations we measure volumes in units that make typical protein concentrations of order unity, so that Ω can be interpreted as a copy number. In accordance with our observations in (Supp. D), a value for Ω can be read as a copy number for Pax6, Nkx2.2 and Irx3; the corresponding typical copy numbers for Olig2 are ten times higher (Supp. D).

However, the model is a coarse-grained description that does not explicitly describe the many possible sources of noise within a living cell. These include spatial heterogeneity and effects from the bursty, multi-step nature of protein production, which includes processes such as transcription, translation, post-translational modification, protein folding and protein shuttling [McAdams and Arkin, 1997]. As noted in [Van Kampen, 2007] and as implemented in [Wang et al., 2007, Zhang et al., 2012, Li and Wang, 2013], Ω relates inversely to the magnitude of fluctuations at a macroscale. It therefore represents the combined effect of all the processes involved in gene regulation and protein production that contribute to the overall system noise. Hence Ω is an “effective” system size parameter, which incorporates all the stochastic effects in the system. Of particular relevance, mRNA molecule number is typically one thousandth that of protein number [Schwanhäusser et al., 2011]. Consistent with this we have found an average of ~100,000 Olig2 protein molecules/cell but only ~40 Olig2 mRNA molecules/cell [Rayon et al., 2019].

We therefore set out to estimate lower and upper bounds on the noise level Ω^−1^, *i.e.* the range of noise that makes sense within our description. The lower bound is given by the typical number of proteins of each species in a cell: these numbers determine the minimum amount of noise that must arise from the stochastic nature of protein production and decay. From protein quantifications (Supp. D) we obtain Ω_max_ ~ 10, 000 for the protein counts of Nkx2.2 and Pax6 per cell at saturation levels (which in our model correspond to concentrations close to unity). Olig2 has a higher estimated count of ~100,000 and in accordance a 10 times higher concentration in the model (the maximum concentration for Olig2 is 10, and 1 for the other TFs). Because of the many neglected sources of additional noise, we expect 1/Ω_max_ to be a considerable underestimate; indeed, simulations with this noise level show almost deterministic behaviour. However, already for a slightly increased noise level (Ω = 2000), we find that the relationships between jump-rate differences across WT and mutant phenotypes discussed in the main text hold true (see Fig. S6). In particular, the WT presents a small amount of heterogeneity (as observed *in vivo*) and the mutants have a more heterogeneous boundary than the WT.

**Figure S6:**
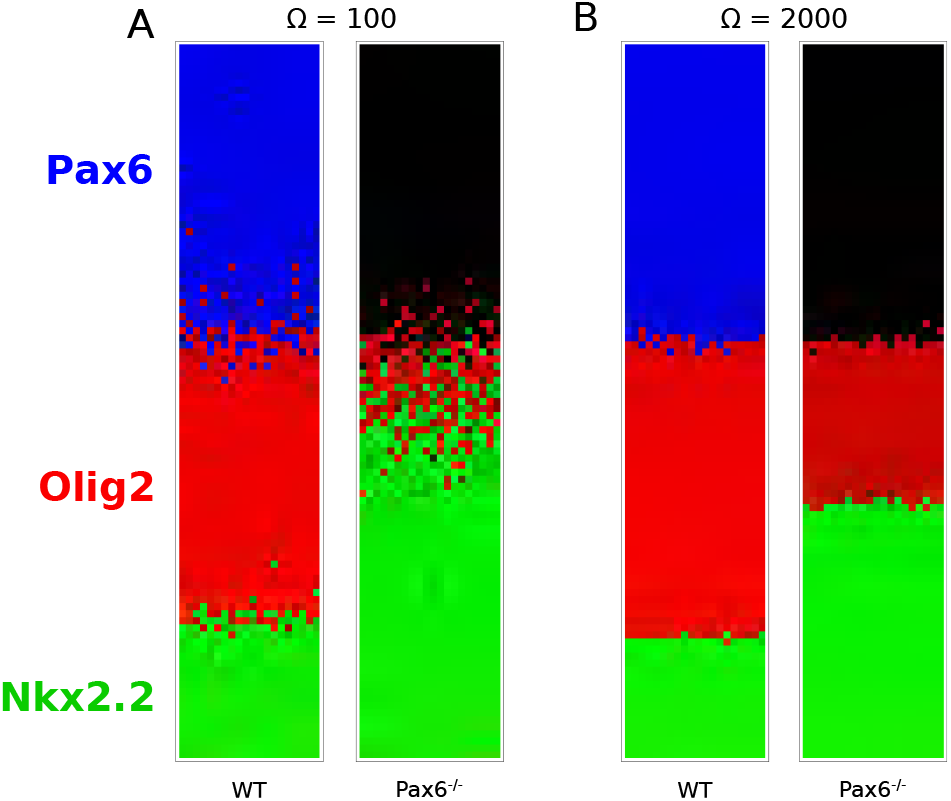
**Simulations of the WT and Pax6**^−/−^ **stochastic models** for (A) Ω = 100, (B) Ω = 2000. For this range for Ω the simulations recapitulate the observed relationship of boundary sharpness and position in WT and Pax6^−/−^ mutants.

To obtain a lower bound for Ω, we measured the coefficient of variation at steady state for all 3 TF values across embryos, to estimate the total amount of noise in the system (Fig. 1A). We then decrease Ω in our numerical simulations until we see coefficients of variation similar to those observed *in vivo*, giving Ω_min_ = 20. This assumes that *all* observed differences in protein levels arise *solely* from the stochasticity in our model. We reason that there are other sources of noise that make the coefficients of variation higher *in vivo*, such as noise resulting from transcription, protein transport within the cell, antibody specificity and measurement error, so that the amount of noise contributed by the stochasticity in our dynamical model will be smaller than 1/Ω_min_ = 1/20. On that basis we find a reasonable smallest value of Ω of ~ 100. The value we use for all results throughout this study is Ω = 250, which is within the broad bounds of Ω_min_ = 20 and Ω_max_ = 20, 000. Importantly, the results we observe remain qualitatively unchanged across the entire range of Ω that we assess as reasonable, 100 ≤ Ω ≤ 2000 (Fig. S6).

To confirm that the effective Ω provides a reasonable estimate of the effect of noise, we performed simulations of the GRN that incorporate the mRNA as well as the protein steps for the production of TFs as additional variables in the system. For this we use experimentally determined mRNA levels [Rayon et al., 2019]. With this addition, protein levels of between 10^4^ and 10^5^ molecules/cell and mRNA levels of ~50 molecules/cell recapitulate the experimentally observed variance in protein levels and the stochasticity of cell fate transitions (Fig. S7). Since it is not our aim to add unnecessary complexity to the model, we did not include mRNA steps in our further analyses.

**Figure S7:**
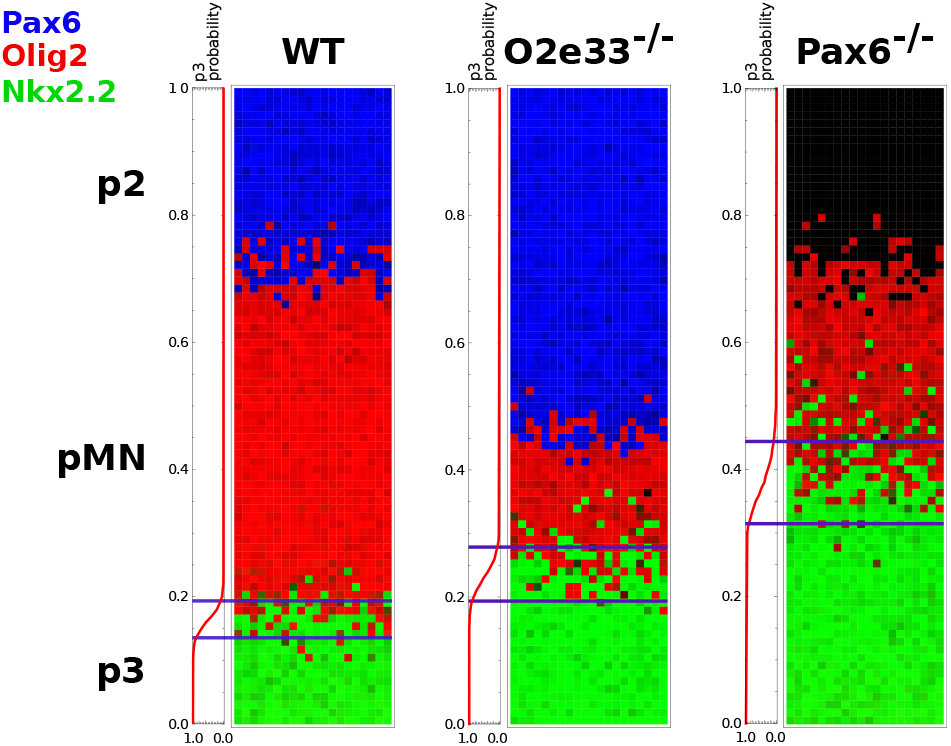
Stochastic simulations of the three genotypes incorporating mRNA as well as protein production. Setup consistent with the simulations shown in the manuscript, with the same colour code and top-down being dorso-ventral. The simulated model includes transcription and translation by including mRNA and protein concentrations for each TF. The simulations use a copy number of 100,000 protein/molecules per cell for Olig2 and 10,000 for the other TFs. For mRNA of Olig2, 40 molecules per cell were used, and 20 molecules for all other mRNA numbers. Parameters for transcription and translation have been extracted from experimental measurements [Rayon et al., 2019]. The lines along each simulation graph show the probability of finding a cell in the p3 state at each neural tube position, with the lines indicating the boundary width extracted from these probabilities.

### Minimum action path

Much of the theoretical analysis in the main text concentrates on the stochastic transitions between fixed points of the deterministic GRN dynamics, which are long-lived metastable states of the stochastic dynamics. The minimum action path (MAP) is the most likely path the system takes in such a transition (for large enough values of Ω), from a steady state to a transition point (which is the saddle point of the dynamical system) and then onwards to a new steady state. The second piece of the path always follows the deterministic dynamics and has a negligible effect on the transition times, so we focus on the first part of the path.

The negative log probability for any path is proportional to what is called the action, which for our Langevin dynamics is of so-called Onsager-Machlup form [Kleinert, 2009]. The action is an integral over time of the Lagrangian, which in turn depends only on the current state (vector of concentrations) and velocity of the system. The time integral can be discretised and the action then minimised as described in e.g. [Bunin et al., 2012]. We analyse the resulting MAP in gene expression space in order to understand how its shape affects the jump times between steady states and thus eventually the boundary width.

The typical time the system takes to reach any point on the MAP scales exponentially with the action up to that point, hence this quantity can be interpreted as an effective energy, within the analogy of a particle making a transition from one local minimum in an energy landscape across a barrier to another minimum. In Fig. 4F we plot this effective energy along the (relative) length of the MAP, describing the effective energy landscape governing the transition. Fig. S8 shows an alternative representation that gives further insight: we plot the derivative of the action along the path, which is the effective force pushing the system back towards the initial steady state.

**Figure S8:**
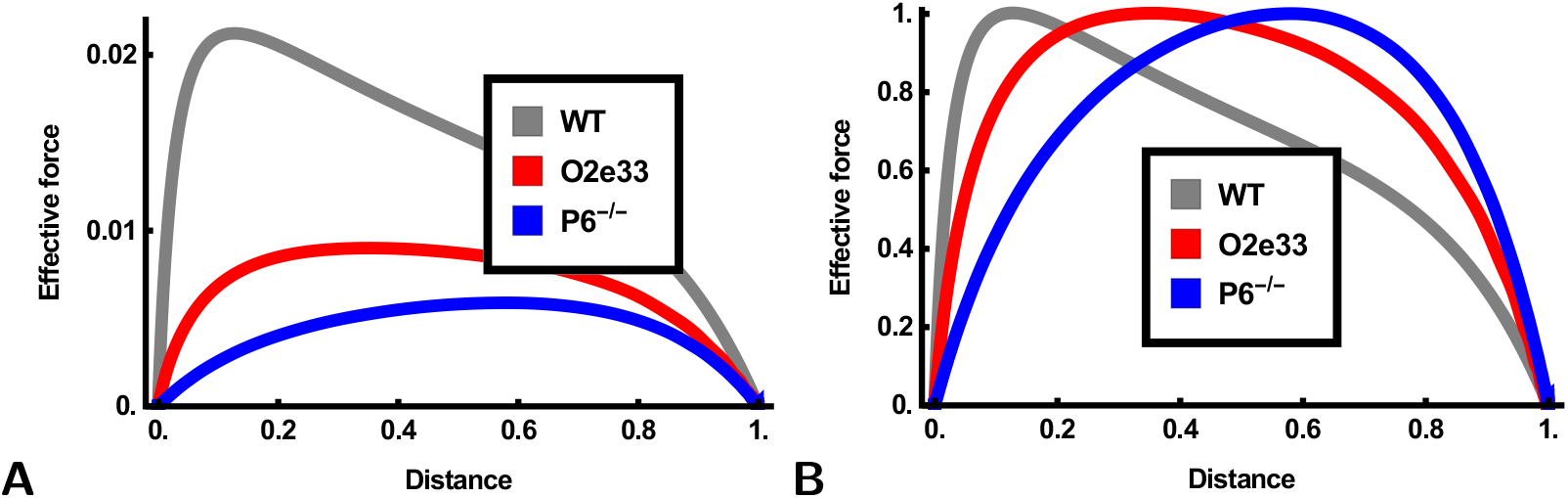
(**A**-**B**) Unnormalised and normalised space derivative of the action along the MAP, plotted along the length of the path. This reflects the effective force driving the system back towards its initial steady state. In the WT system (gray) the force is highest near the beginning of the path, leading to a noticeably skewed plot, while the O2e33^−/−^ (red) and Pax6^−/−^ (blue) more nearly symmetric force profiles. The high initial force in WT responsible for the large typical jump times in the system, and is related to the significant curvature of the MAP away from the straight line between initial steady state and transition point (Fig. 4D-F)

### Calculating magnitude of fluctuations

To compare the magnitude of fluctuations between WT and mutants *in silico* we take two separate approaches. The first is to consider fluctuations in expression levels around a steady state, before any transition to a new state occurs. For moderate noise levels such fluctuations can be analysed using a linear expansion of the dynamics around the steady state (here: pMN), leading to a local Gaussian distribution of expression levels. The corresponding covariance matrix ***C*** can be calculated from the Jacobian matrix ***J*** of the linearized dynamics and the noise covariance ***D*** as defined in (C.1b), both evaluated at the steady state. The required link between the three matrices is the Lyapunov equation, which determines ***C*** via

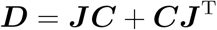

Once ***C*** has been found we normalise it by the corresponding pMN steady state values (***X***), to obtain 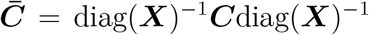. We finally compute the trace of 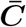 and take the square root. The end result is the typical standard deviation (root-mean-square fluctuation) of the expression levels, relative to the mean expression levels. This is shown in Fig. S9**A** as a function of neural tube position.

**Figure S9:**
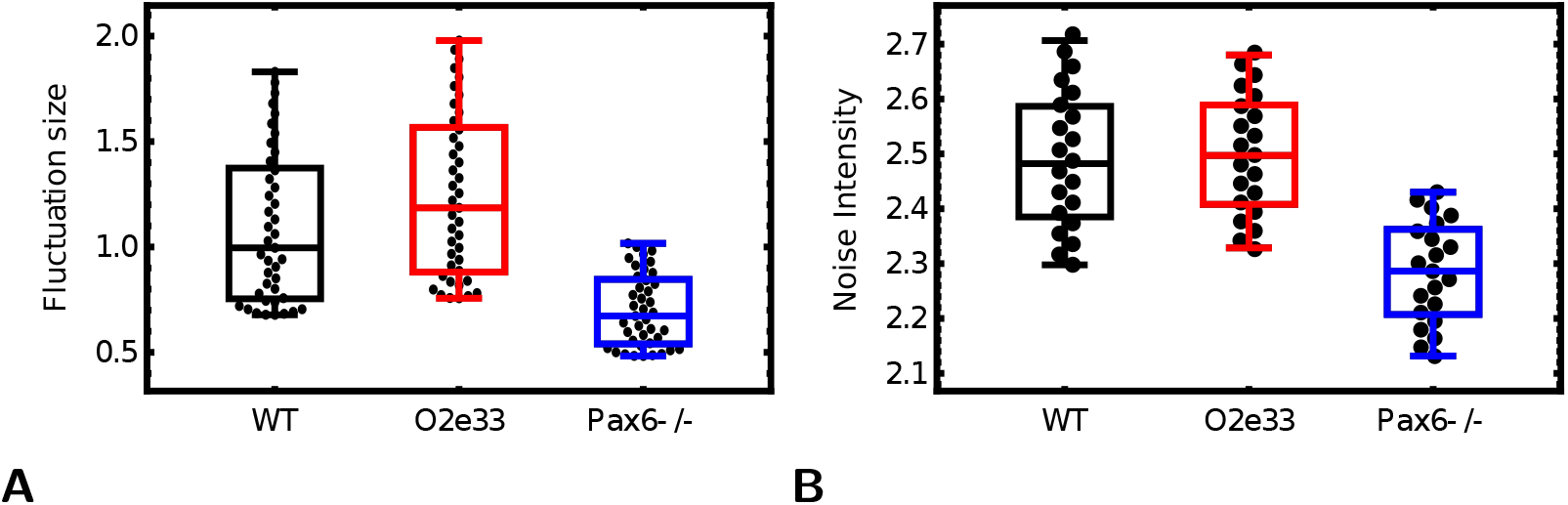
Comparing total noise across genotypes. (A) Comparison of noise levels as defined by root-mean-square relative expression level fluctuations, calculated within a Gaussian approximation near the steady state. Points represent different positions along the neural tube (B) Noise levels defined as noise variance calculated at equidistant points along the MAP, at fixed fractional neural tube length from the bifurcation point. Note that in both definitions, noise levels are comparable across WT and both mutants, with slightly lower values in Pax6^−/−^.

The second approach to quantifying noise levels is to use the noise variance, which is the trace of the noise covariance matrix given in (C.1b). This noise variance depends on the expression levels so we average it across equidistant points along the MAP and take the square root of this value to obtain the root-mean-square noise level. Example results at a specific position along the neural tube are shown in Fig. S9B; results at other positions were qualitatively the same (data not shown). Both approaches to quantifying noise show comparable total variance across the different genotypes, with slightly lower noise in Pax6^−/−^ than in WT and O2e33^−/−^. To make the comparison to *in vivo* observations we accounted for the fact that experimentally, noise levels are averaged across several neural tube positions throughout the pMN domain. We therefore also performed an average *in silico* of neural tube positions to obtain comparable data for Fig. 3**D**.

## D Protein Number Quantifications

**Figure S10:**
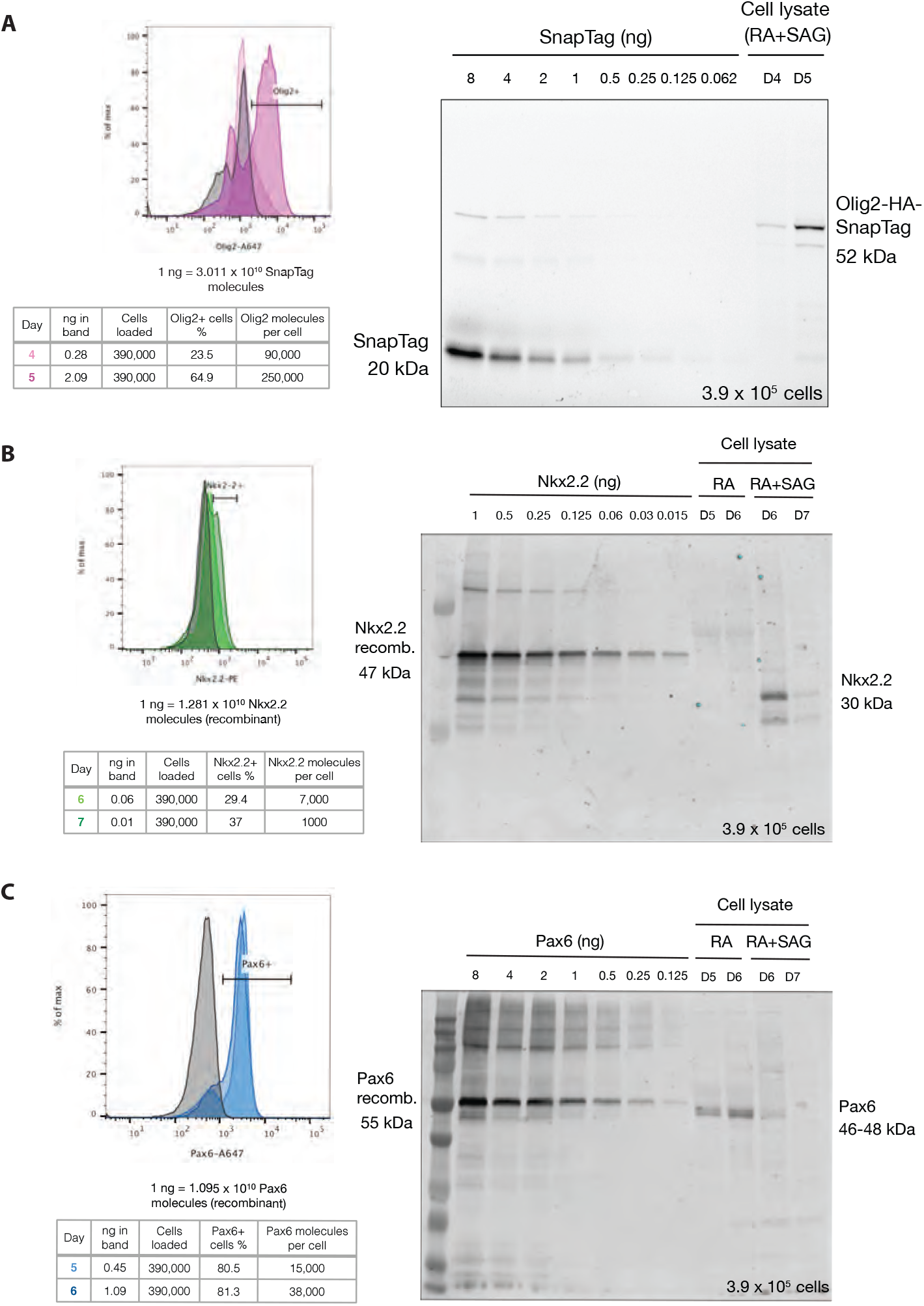
Quantifying Protein Copy Number. (**A**) Flow cytometry analysis to determine percentage of Olig2 expressing cells in differentiated ES cells at the indicated days. Table shows quantification of a gel for days 4 and 5. Olig2 has approximately a 10-fold higher protein copy number compared to Nkx2.2 and Pax6. (**B**) Analysis of Nkx2.2 expressing cells on days 6 and 7 of differentiation. Nkx2.2 molecules per cell calculated using the measured percentage of cells expressing Nkx2.2 and quantification of the Western blot analysis. (**C**) Analysis of Pax6 expressing cells to determine protein copy number at days 5 and 6 of differentiation. Pax6 molecules per cell calculated using the measured percentage of cells expressing Pax6 and quantification of the Western blot analysis.

## E Simulating WT and mutant GRNs

We used the equations and parameters described in [Cohen et al., 2014] for the GRN that patterns the neural tube; this parameter set was optimised to replicate the boundary positions in wild-type and mutant embryos. Following the inclusion of the noise term as explained in Supp. C we explored the effect of the initial conditions for the TFs (i.e. their initial expression levels *x*_*j*_). The aim was to find a consistent set of initial conditions that sustain the boundary positions but also recapitulate the boundary sharpness of each mutant. The initial conditions that satisfied these conditions were identified in a systematic scan as *x*_Pax6_ = 0.1, *x*_Olig2_ = 0, *x*_Nkx2.2_ = 0, *x*_Irx3_ = 0.1. The p3-pMN boundaries in WT, Irx3^−/−^, Nkx2.2^−/−^ and Olig2^−/−^ simulations remained sharp as is the case *in vivo* (Fig. S11). Only the loss of Pax6 resulted in decreased boundary sharpness. Boundary positions remained consistent with *in vivo* observations as was the case in the original deterministic model (Fig. S11) & [Cohen et al., 2014].

**Figure S11:**
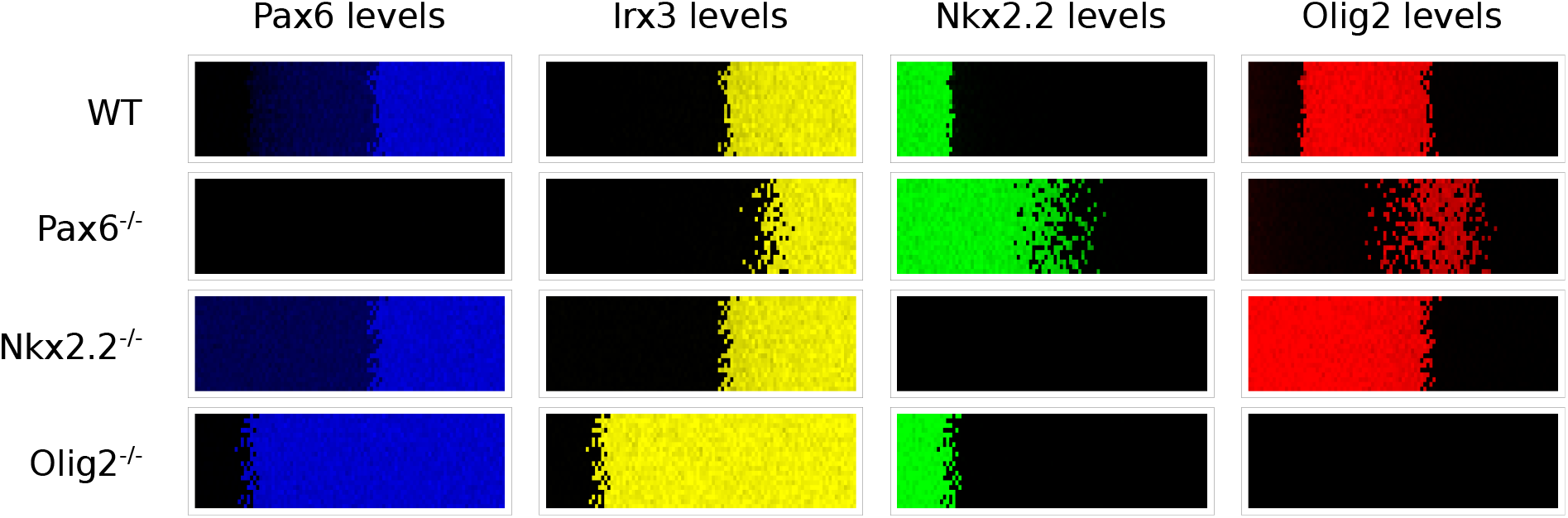
Patterning phenotypes produced by stochastic simulations for WT and mutants. Predicted expression patterns for the four TFs in the indicated genotypes are qualitatively similar to those in [Cohen et al., 2014]. Ventral to the left and dorsal to the right. Although boundary positions change, boundary precision is largely unaffected except for Pax6^−/−^, consistent with *in vivo* experimental observations.

### Model parameters

We detail the parameters used throughout the paper to model neural tube development for equation (C.1a), and adapted for the computational screen as explained in Supp. F.

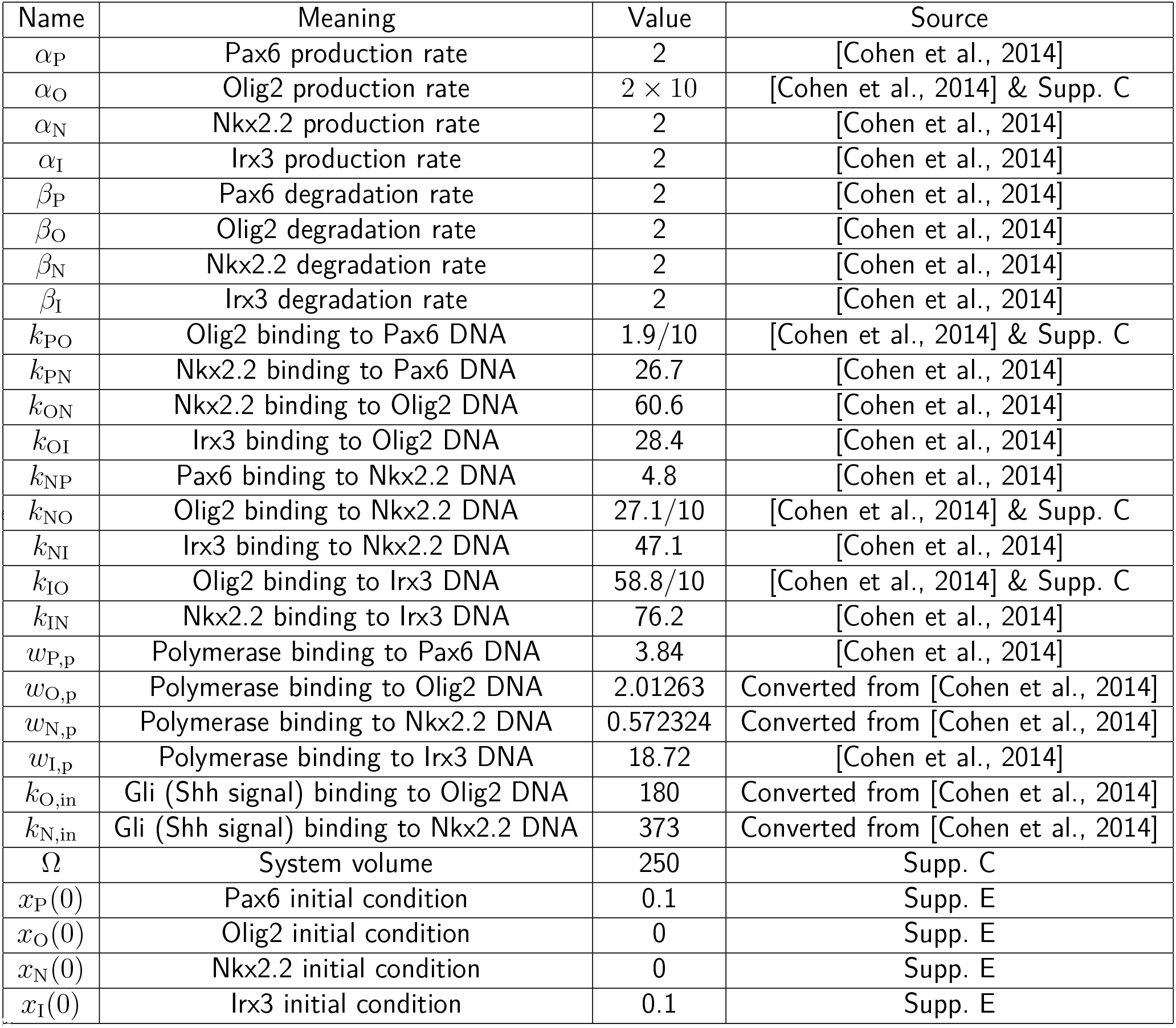

Where factors of 10 have been written in the table, these arise because we have modified the model of [Cohen et al., 2014] to represent explicitly the experimental observation that Olig2 has a concentration 10 times higher than the other TFs. While this difference is immaterial for a deterministic description of the GRN dynamics, it affects the stochastic representation because larger copy numbers have smaller relative fluctuations.

The above parameters are used in the general model (C.1a) for the dynamics of the TFs *j* = P (Pax6), O (Olig2), N (Nkx2.2) and I (Irx3). DNA conformations are defined by the numbers ***n*** = (*n*_p_, *n*_in_, *n*_P_, *n*_O_, *n*_N_, *n*_I_) of bound molecules of polymerase, Gli signal input, Pax6, Olig2, Nkx2.2, Irx3 in that order. The only allowed conformations are the empty conformation, the conformations with polymerase and *n*_in_ = 0 or 1 signal molecule bound; and conformations with at least one molecule of the other TFs bound, with maximally two molecules from each other TF. All other conformations are assigned affinity zero. The weights for the allowed conformations are multiplicative, with bound polymerase contributing a factor *w*_*j*,p_ (see below), bound signal a factor *k*_*j*,in_*x*_in_ and each TF *i* bound to DNA producing TF *j* a factor *k*_*ji*_*x*_*i*_. Examples of the corresponding affinities are *k*_O,(0,0,0,0,1,0)_ = *k*_ON_ and *k*_O,(0,0,0,0,0,2)_ = *k*_OI_^2^. The polymerase binding parameters are directly stated as the weights *w*_*j*,p_ = *k*_*j*,p_*x*_p_ including polymerase concentration (which is assumed constant). As detailed in [Cohen et al., 2014], this weight describes all basal production inputs for each TF and thus represents input from TFs such as Sox2 [Graham et al., 2003, Oosterveen et al., 2012, Peterson et al., 2012]. Finally, the protein production rates *α*_*j*,***n***_ in the general model (C.1a) are set to the value given in the table for the DNA conformations with bound polymerase, and zero otherwise.

As an explicit example of the resulting GRN equations, we write here the production rate for Olig2:

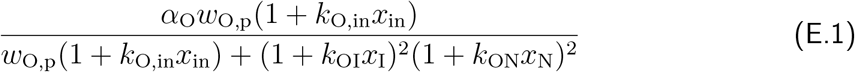

The signal input concentration *x*_in_ is the gradient *e*^−*s*/0.15^, which depends on the dorsal-ventral neural tube position *s* ranging from 0 to 1 as in [Cohen et al., 2014].

### O2e33^−/−^ mutant

To find parameter sets that describe the behaviour of the O2e33^−/−^ enhancer mutation, we first identified those parameters that are related directly to the deletion of the respective enhancer. Analysis of the sequence of the enhancer together with CHIP-seq and ATAC-seq [Oosterveen et al., 2012, Peterson et al., 2012, Kutejova et al., 2016, Metzis et al., 2018] suggested that Gli proteins, Nkx2.2, Irx3, and Sox2 all have a direct effect on this enhancer (Fig. 2A). We therefore considered variations in the parameters that specify Nkx2.2 binding, Irx3 binding, Gli binding and basal production (corresponding to Sox2 binding). We systematically explored how reducing the parameters for each of these interactions, to a fraction *f* of their original value, could explain the observed phenotype. We used a uniform distribution to perform this search and throughout this supplementary represent the respective parameter reductions directly in terms of the ratio *f* between new and original (WT) parameter values.

### Fitting *in vitro* delay and resulting predictions

We first identified parameter sets that could replicate the observed *in vitro* delay in the onset of Olig2 expression in the mutant, leading to a reduced parameter space (Fig. S12). In this step we do not set any constraints to the position or precision of boundaries between expression domains as this information cannot be extracted from the *in vitro* system. The delay in Olig2 activation was determined for networks positioned a fraction 0.3 along the neural tube, and we retained those networks that took twice the amount of time to express Olig2 than in the WT. The same measurement was performed at other neural tube positions and resulted in similar distributions (data not shown).

We next investigated what further phenotypical behaviour the retained parameter sets predict, focussing on the domain size and boundary precision generated in response to a graded Shh signal. We found that 68% of the parameter combinations reduced boundary precision, 80% reduced the size of the pMN domain, with 83% presenting one or the other of the phenotypes (data not shown). Here, the pMN domain size was calculated with respect to the Shh gradient and we considered it reduced if it was below 70% of the WT size. For determining boundary sharpness, we regarded as imprecise those systems that had a boundary width at least twice the size of the WT; this width is calculated using the SDE system with the thresholds described in Fig. 3B. The fact that a majority of the parameter sets identified affected domain size and boundary precision encouraged us to generate the mouse lines.

**Figure S12:**
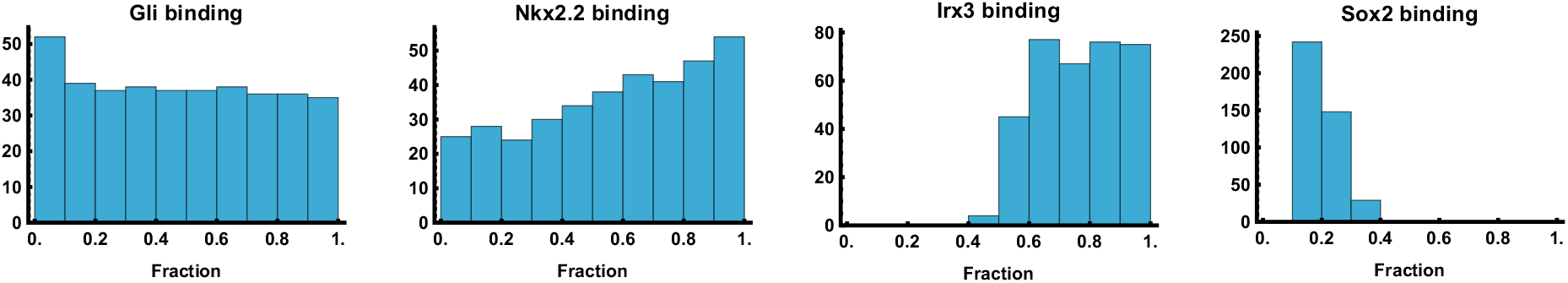
Distribution of parameter changes to mimic *in vitro* O2e33^−/−^ mutant. To recapitulate the O2e33^−/−^ dynamics *in vitro*, model parameters were systematically explored to identify changes that could account for the delay in onset of Olig2 expression. The graphs show the distributions of reduction factors *f* (x-axis) relative to WT parameter values, across parameter sets that recapitulate the delay. The (y-axis) shows number of parameter combinations that recapitulate the phenotype. The results show that what is needed to generate a delayed induction of Olig2 is a substantial reduction in Sox2 input while maintaining input of Irx3.

### Fitting *in vivo* phenotype with patterning information

Once the mouse lines were generated we confirmed the delay in onset of Olig2, and noted two additional phenotypes as expected from the initial parameter screen: a loss of precision at the p3-pMN boundary and a ventral shift of the pMN-p2 boundary. Importantly, this *in vivo* data allowed us to define targets regarding boundary position and precision for our fitting of the mutant phenotype. The new targets were therefore extracted from this data, and were used to further constrain the results displayed in Fig. S12. These additional constraints were:

- The pMN-p3 boundary width to be at least twice the size of the WT as explained above.
- The pMN-p3 boundary position to be between [0.17 0.25] (as the WT boundary position is at 0.17 and some of the *in vivo* mutants show a small dorsal shift).
- The p2-pMN position to be below or equal to 0.5 (WT boundary is at 0.55, this means a reduction of the domain size of at least 15% with respect to WT) but higher than the pMN-p3 boundary position, such that the pMN domain does not disappear.
- Other aspects of patterning not to be disturbed.

The resulting retained networks present a substantially reduced parameter space and are shown in Fig. S13. From these parameter sets we took a representative point as our model for the O2e33^−/−^ mutant; as expected this replicates the observed experimental phenotypes.

**Figure S13:**
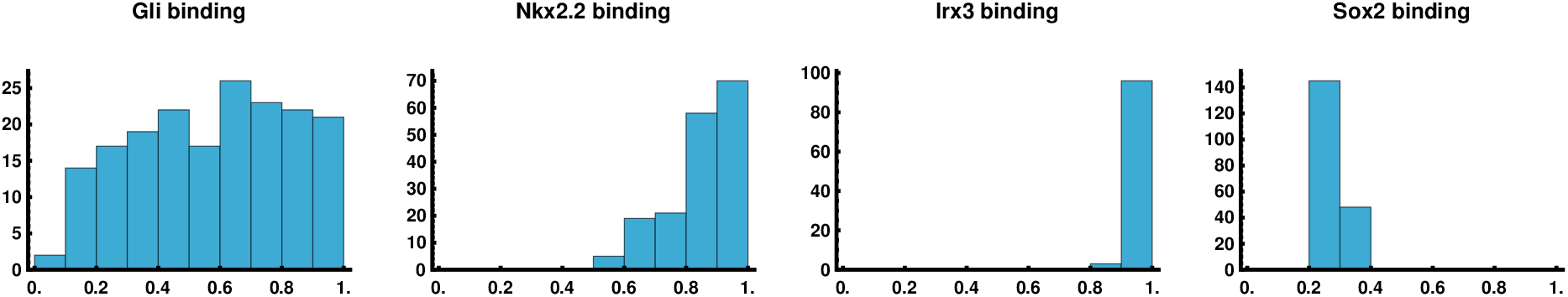
Distribution of parameter changes to mimic *in vivo* O2e33^−/−^ mutant. Equivalent histograms to Fig. S12 with the additional constraints from *in vivo* observations: ventral shift of pMN-p2 boundary *and* broad p3-pMN boundary. The main results are: maintaining WT levels of Irx3 input; substantial reduction in Sox2 input, some reduction in Gli input but with a broad distribution, and a mild reduction in Nkx2.2 input.

## F Screening three node networks for precision

### Defining a functional form

To perform a parameter screen we explored three node networks with all possible interactions between the nodes, as this has provided useful insights in other systems (Fig. 1A) [Cotterell and Sharpe, 2010, Leon et al., 2016]. For the purpose of exploring different dynamics, we enumerated the different possible transcriptional/occupancy states of the promoter to model the production rates of a given protein. These rates depend on polymerase availability, signal input (morphogen) and regulating transcription factors, with concentrations *x*_p_, *x*_in_ and *x*_*i*_ respectively. The transcription factors *i* can be activating 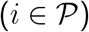 or repressing 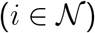, with 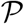 and 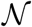 denoting the sets of activating and repressing transcription factors, respectively. While in the previous model, in its most general form (C.1a), different protein production rates can be used for different DNA conformations, in the neural tube network we used the same production rate for all protein-producing input conformations (see Supp. E). We adopt the same approach here and set the production rate to unity in appropriate units of time; thus the model is specified only by the binding affinities of the various DNA conformations. Without loss of generality we fixed the affinity (and hence the weight) of the unbound conformation to 1 as explained in [Sherman and Cohen, 2012]. We assign the weights of conformations with only one bound molecule as *k*_p_*x*_p_, *k*_in_*x*_in_ and *k_i_x_i_*. In accordance with our previous model (C.1a), we set the following constraints:

- All conformations with polymerase and without any repressor 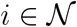 produce protein; it does not matter whether signal or any activator 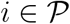 are bound.
- Conformations that have one or more repressor 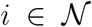 bound together with either signal, polymerase or any activator 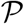 are excluded, based on the assumption that these molecules compete for the same binding site
- Binding of signal or any activator 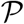 enhances binding of polymerase
- No other cooperativity effects are present

### Expressions for conformation states

The only states that can produce protein are those with polymerase bound. For brevity we follow the convention in Supp. E and abbreviate

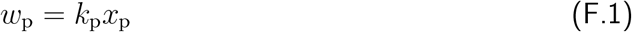

in the following, taking polymerase levels as constant throughout our dynamics. As specified above, the only states that can bind polymerase are those that have no repressors bound. We assume no cooperativity between signal *x*_in_ and activators *x*_*i*_, 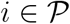, hence the total weight of states that can potentially bind polymerase (assuming two binding sites per activator 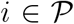 but only one for the signal) is:

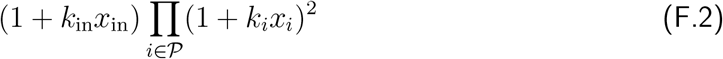

Given that repressors 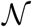 can only bind by themselves, and that there is no other cooperativity between the inputs, the total weight for conformations with at least one repressor 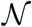 bound while assuming two binding sites per repressor 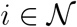 is:

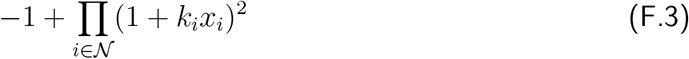

In accordance with biological intuition, polymerase is recruited by activators 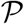 or signal. The simplest way to implement this is to increase the weight of conformations having both polymerase and at least one activator 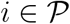 or signal by a cooperativity factor *c*, giving a total weight of:

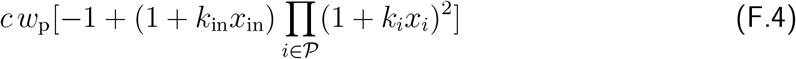

Finally, the weight for the unbound (empty) conformation is taken as 1, as explained above, and for the conformation with one polymerase bound it is *w*_p_ as defined in (F.1). The total weight, i.e. the denominator of the protein production rate, is then

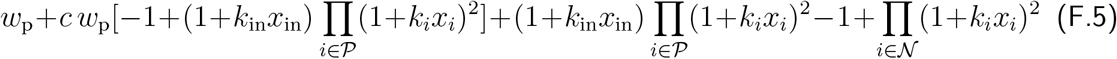

while the numerator is the total weight of conformations *with* polymerase, either on its own (F.1) or together with activators or signal (F.4), giving overall for the production rate (which with protein production set to unity is also the probability of being in a DNA conformation that produces protein)

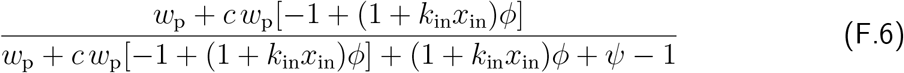

with the abbreviations

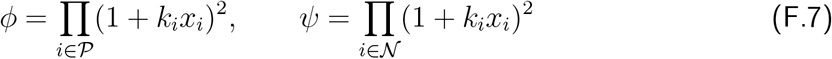

### General strong cooperativity limit

It will be convenient in the following to write the effective affinities of signal and activating TFs in combination with polymerase in a form that includes the cooperativity effect from the factor *c*, i.e. in terms of 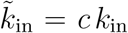 and 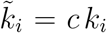 for 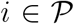. The protein production rate is then expressed as

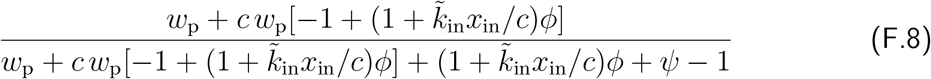

with now

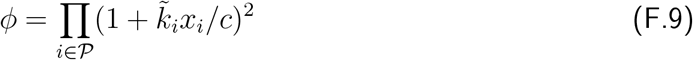

We can now compare with the analogous expression (E.1) in the neural tube network. There all interactions are repressive so that 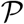 is the empty set and hence *ϕ* = 1, which simplifies (F.8) to

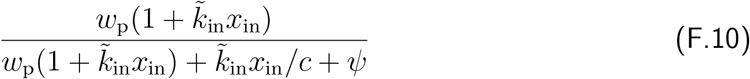

This agrees with (E.1) except for the middle term in the denominator, which represents the weight of DNA conformations with only signal but no polymerase bound. Its absence in the neural tube network formally corresponds to the strong cooperativity limit *c* → ∞. In our screen we use a finite cooperativity *c* = 100 to avoid the extreme case of excluding conformations with only signal bound completely; this value of *c* is still large enough, however, to replicate the dynamics of the neural tube network. We thus take (F.8) with *c* = 100 as the form of protein production rates in our screen; compared to the neural tube case this allows us to include both activating and repressive interactions.

Adding a protein decay term (with unit decay rate) and stochastic fluctuations, the dynamics of the three-node networks in our screen, with protein levels *x*_1_, *x*_2_ and *x*_3_, is thus described by

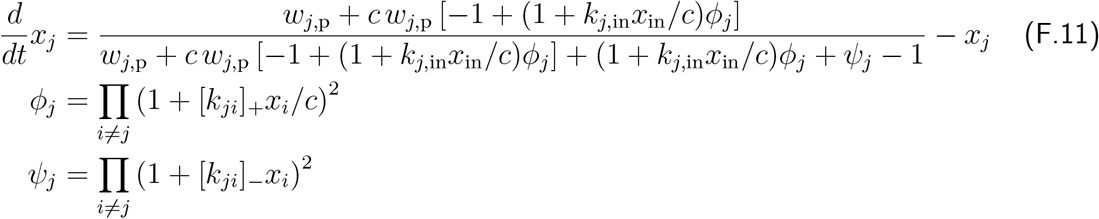

for *j* = 1, 2, 3; compared to (F.8) we have dropped all tildes to unclutter the notation. We have also allowed the sets 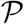 and 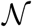 of activating and repressing transcription factors to be determined implicitly by the system parameters. This is done by generalizing the affinities *k*_*ji*_ so that a positive sign indicates an activation of *j* by *i* and a negative sign a repression. The corresponding switching of species *i* between the products over activators and repressors is achieved mathematically by setting [*k*]_+_ = max(*k,* 0) and [*k*]_−_ = max(−*k,* 0).

To mimic the structure of the neural tube network, we assume that only proteins 1 and 2 have direct signal inputs, while 3 does not, so that *k*_3,in_ = 0. This leaves 11 network parameters: 2 for the signal (gradient) inputs from the gradient (*k*_1,in_ into node 1 and *k*_2,in_ into node 2), 6 from the interactions between TFs (*k*_12_, *k*_13_, *k*_21_, *k*_23_, *k*_31_ and *k*_32_) and 3 for polymerase binding weights (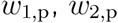 and 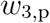).

### Parameter exploration

We explored the 11 dimensional parameter space specified above using a uniform log distribution (log_10_), where the ranges are set differently depending on the parameter. Specifically we chose the ranges as: range(*k*_in_) = [10 : 400], range(*w*_p_) = [0.1 : 10], range(*k*_*ji*_) = [−100 : −1] ∪ [1 : 100] with the sign of each regulation *k*_*ji*_ being chosen randomly.

We provide a schematic in Fig. S14 of the sequential steps taken to screen for relevant networks, analyse them and classify them into topologies. We explored parameter combinations for a three node network defined in the form (F.11). The main criterion for choosing a viable set of parameters was that they must produce a patterned steady state, i.e. a saddle-node bifurcation on the same gradient as in the neural tube, defined as *x*_in_ = *e*^−*s*/0.15^ where *s* indicates dorsal-ventral neural tube position and ranges from 0 to 1. To avoid trivial effects from shifts in the boundary position we set a further constraint that the bifurcation must occur at a position *s* in the same range as in the neural tube network, 0.165 ≤ *s* ≤ 0.17. More specifically networks were required to be monostable below *s* = 0.165, with high levels of *x*_1_; and bistable beyond *s* = 0.17, with one state having high *x*_2_ and the other high *x*_1_ (with “high” being a concentration value above 0.6). For each network meeting these criteria, we then proceeded to calculate the MAPs in the same way as for the neural tube network (as explained in Supp. C), and the jump time. We selected networks that have boundaries sharper than a certain threshold, set by requiring the boundary to be no wider than 0.2 fractional neural tube units; boundary widths were calculated based on their transition time obtained from simulating the SDEs. To simulate the neural tube network from (E.1) in the screen we used the standard parameters from that network, reverting to the original version [Cohen et al., 2014] with maximal concentrations of unity for all TFs in order to ensure comparability with the networks produced by the screen. We removed all terms relating to Irx3, as these do not contribute substantially to the dynamics of transitioning from a pMN to a p3 steady state. We further set production and degradation rates to be equal to unity in the screen as these simply scale the jump time and do not affect the results.

**Figure S14:**
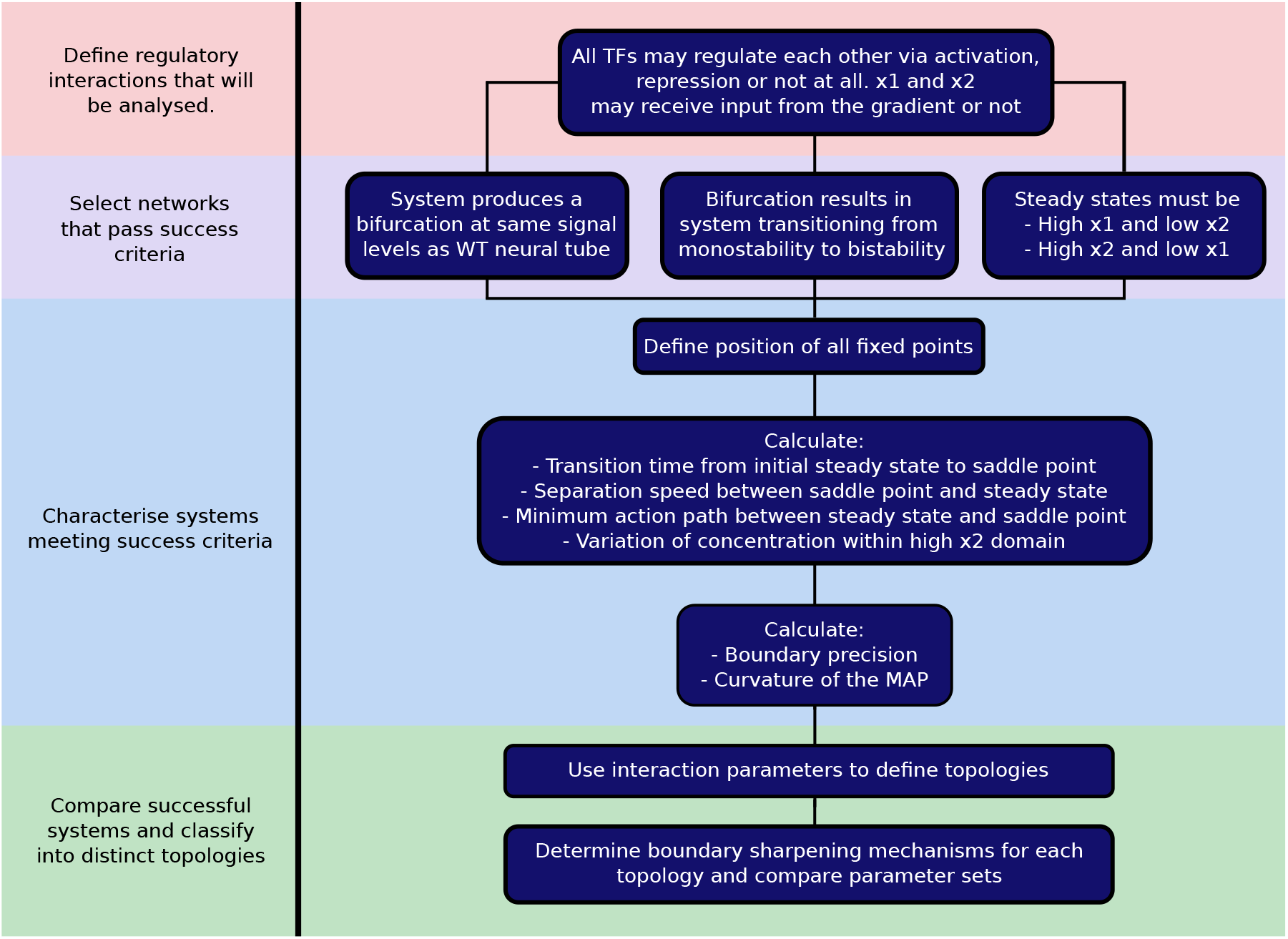
Schematic of steps for systematic screening. We desgined the screen to first identify parameter sets that describe networks that generate a sharp boundary at a specific location within a gradient. This ensures that the resulting networks are comparable with each other. The parameter sets that pass this filter are then analysed by defining the characteristics relevant to forming a precise boundary. Finally, we classify the parameter sets into topologies.

In analysing the results of the network screen we quantified the curvature of the MAP as the largest perpendicular distance of any point on the MAP from the straight line between steady state and transition point, normalised by the total length of this line. We refer to this value throughout the text by the shorthand “curvature” as it gives a quantitative indication of how much the MAP deviates from the shortest path. The curvature was measured at *s* = 0.25 and the robustness of the results with respect to this choice of neural tube position was tested by comparing with multiple other locations, with qualitatively similar results in all cases (data not shown).

In the analysis we also characterised networks by the strength of the contribution of the third node, which does not receive direct signal input. We quantified this by taking the value of *x*_3_ at the steady state and transition point (saddle point) and multiplying each by parameters for the repression or activation of nodes 1 and 2 by node 3, taking the maximum value. The multiplication by representative concentration levels of the third node was motivated by the fact that when those concentrations are small, even large interaction parameter values have small net effects.

Networks with a low third node contribution are effectively two node networks, and turned out to have low MAP curvature. This led us to explore other mechanisms for generating sharp boundaries. Geometrically, in the space of expression levels (phase space), the speed at which the steady state and saddle point separate as a function of neural tube position *s* is a plausible contributor to boundary sharpness because even if the fluctuations around the initial steady state favour a jump, such a jump will be inhibited by a large separation between steady state and transition point. High separation speed should thus lead to rapidly increasing jump times and hence to sharp boundaries. To measure separation speed we focussed on a fixed position (chosen as *s* = 0.25) along the neural tube, beyond the saddle-node bifurcation, and calculated the Euclidean distance between steady state and transition point. We then used this as a simple quantitative indication of separation speed. We checked the robustness of this measure by performing the measurement for different *fixed* positions along the neural tube, and also at *variable* locations chosen as the centre of the boundary region for each network; we found qualitatively similar results in every case (data not shown).

**Figure S15:**
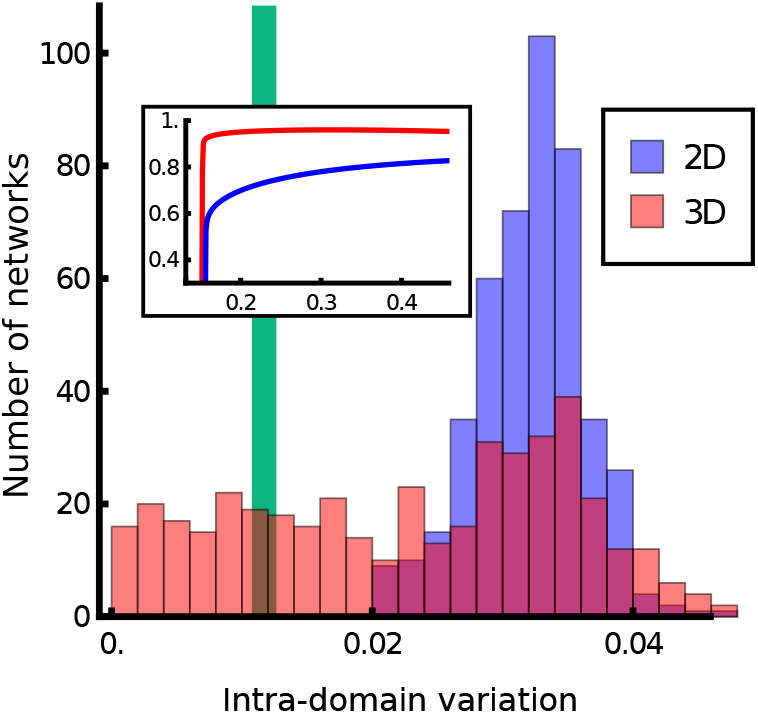
Histogram of variation of the expression level of the second node within its domain of expression for 3D (red) and 2D (blue) networks; inset shows example variation of expression levels across a domain. 3D networks can generate domains of expression with more constant levels of expression (lower domain variation) than 2D networks, which rely on separation speed to create sharp boundaries. Green line represents the WT network.

When a network had a high separation speed, this typically resulted in the steady state (the expression profile) of *x*_2_ varying, i.e. changing within a domain of the steady state pattern.

We quantified this heterogeneity by the standard deviation of *x*_2_ within the region of high *x*_2_ expression. This confirmed (see Fig. S15) that sharp 2D networks have a higher level of heterogeneity than 3D networks, which use the curvature of the MAP to generate sharpness.

### Characterisation of topologies

Finally we analysed the topologies of the networks resulting from the screen. To sort networks into topologies we used thresholds to identify whether nodes 1 and 2 receive significant signal input, and for each of the TF nodes whether it significantly activates or represses the other TFs. Starting with the former, within the input parameter range [10 : 400] for nodes 1 and 2, we took any parameter 30 < *k*_in_ to be a positive input; lower values were classified as lack of input. This cutoff was chosen by testing a range of different values and imposing the constraints that we want to neither classify the majority of networks as having two inputs (which would provide no information on the input topology, as could happen if the cutoff was too low) nor assign any network to a topology with no inputs (which would not make biological sense and would occur when the cutoff is too high). For interactions between nodes we took into account not only the parameters *k*_*ji*_ but whether each parameter in conjunction with the actual states of the system would have a noticeable effect. We evaluated interactions by considering the contribution of an interaction given the highest level that the effector node can take. Accordingly, we consider an interaction with 0.3 < |*k*_*ji*_| max(*x*_*i*_) to be significant, otherwise we classify it as negligible. The maximum was taken over all steady states for all neural tube positions. The cutoff value of 0.3 was chosen by systematic inspection of a representative number of networks, for which we compared the dynamics with and without individual interactions and assessed whether these were qualitatively identical or not. To assess the robustness of the cutoff value, we varied it within a range up to an order of magnitude larger and found that the results of our characterisation of network topologies remained qualitatively the same (data not shown).

With this approach we classified all the 3D network parameter sets into topologies, determined those that occurred most often (Fig. S16) and plotted the boundary precisions they generate (Fig. 5H). The results indicated that although some topologies are more frequently represented amongst networks producing a sharp boundary, there is no single topology that ensures sharpness. Some networks (such as 1–4 in Fig. S16) prevented the boundary from becoming very imprecise, but even within these network topologies the range of sharpness was large (Fig. 5G,H & Fig. S16 & Fig. S17). This leads to the conclusion that the dynamical properties generated by the network, rather than the structure of the network determines boundary precision. Indeed, we confirmed by analysing each topology separately that the main indicators of sharpness are the two mechanisms identified in the main text: curvature of transition path and separation speed (Fig. S17). Nonetheless, a network’s topology can substantially bias the dynamics towards high MAP curvature, and hence towards sharpness.

**Figure S16:**
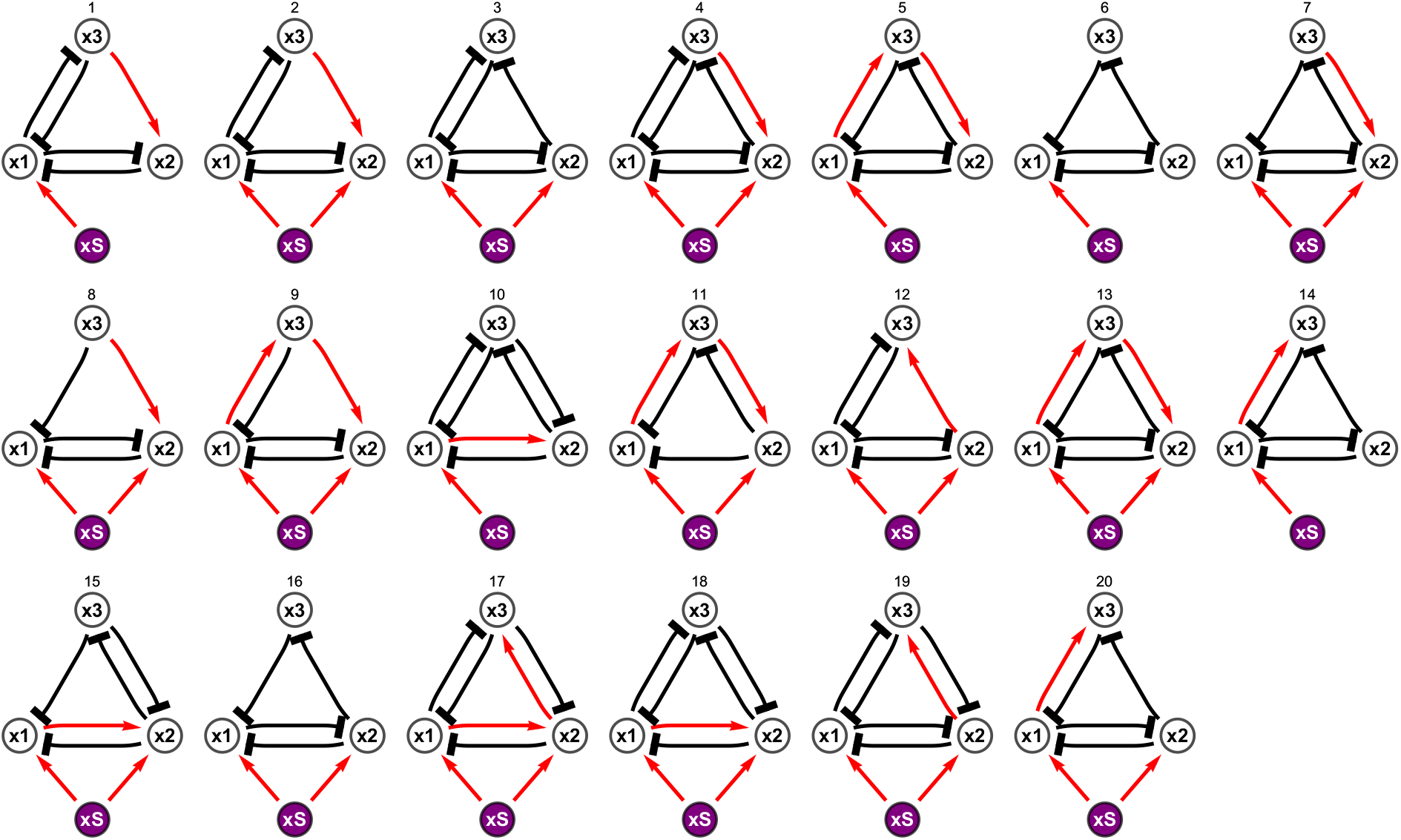
List of topologies that generate sharp boundaries, sorted in the same order as Fig. 5**H**. Red arrows indicate activation, black lines with blunt ends represent repression. Mutual repression between the first and second nodes (1 and 2) is a consistent feature, as well as the input from the signal to the first node. For the sharpest networks, a mutual repression between the first and third nodes is observed.

**Figure S17:**
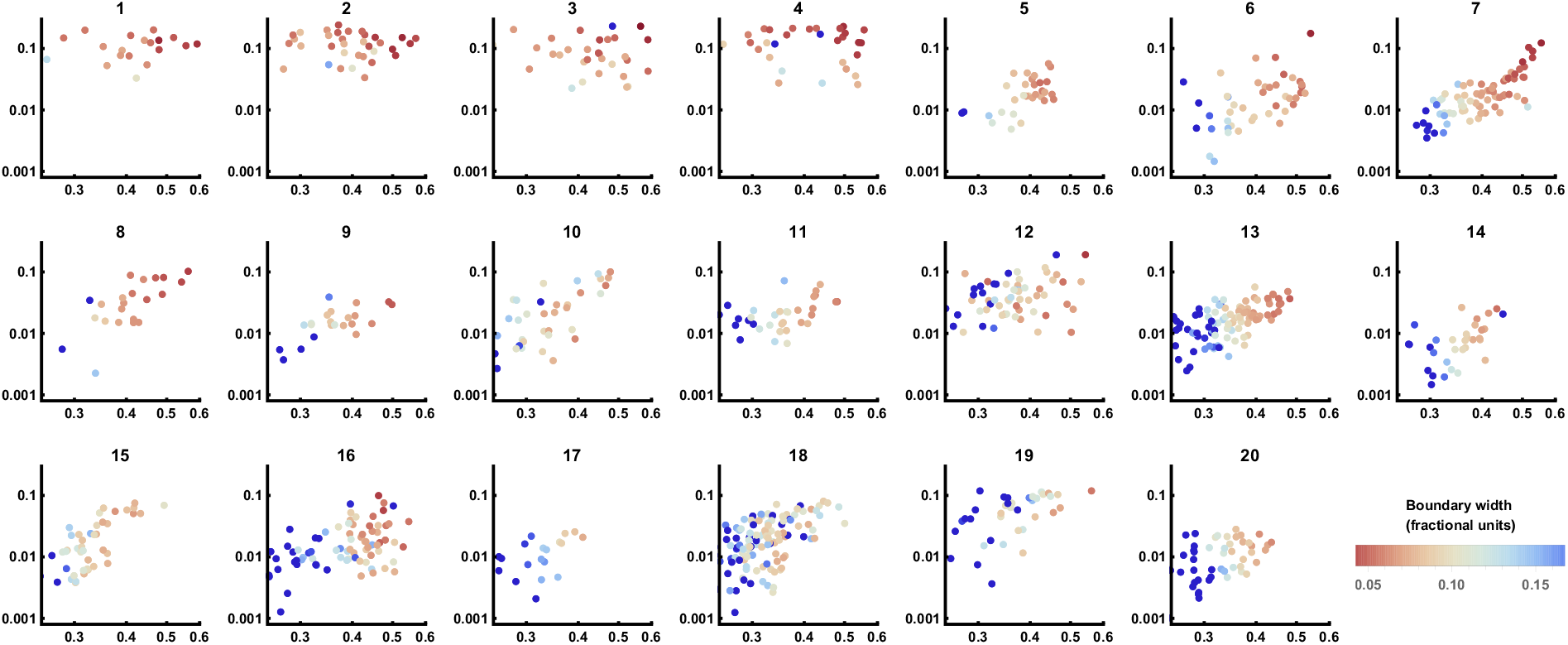
MAP curvature plotted against separation speed with boundary width indicated by colouring. The data are equivalent to those shown in Fig. 5**D**, but here each plot represents an individual network topology and networks with wide boundaries have been included in the plots (deep blue). Network topologiess are ordered as in Fig. S16. While separation speed does not exhibit obvious differences between topologies, network topologies 1–4 have consistent high curvature.

### Effect of signalling noise on boundary precision

We explored what effect noise in the signal gradient would have on the precision of boundaries generated by the mechanisms revealed in the screen. To this end, we simulated networks recovered from the screen using a noisy signal as an input. For this we have used Ornstein-Uhlenbeck noise and explored systematically a range of fluctuation timescales and noise amplitudes (see Eq. F.12). As is commonly done we use a log version to avoid negative values, i.e. we write the fluctuating signal input as *s*_OU_(*t*) = exp(*l*(*t*)) where *l*(*t*) evolves in time as

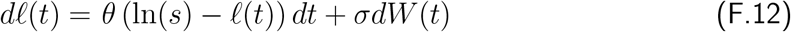

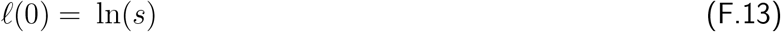

The variables are the standard terms for Ornstein-Uhlenbeck processes: *θ* is the inverse correlation time of fluctuations, *σ* is the noise amplitude, *W* is a Wiener process, *s* is the constant Gli input in the original model. We compared the boundary widths generated by simulations using these noisy gradients with those in which the signal was constant, for otherwise identical parameter sets (Fig. S18). This revealed that noise in the signal had relatively limited effects on the precision of boundaries for moderate levels of noise. Moreover, the same relative sharpness of boundaries for the different networks was found in the simulations with a constant and a noisy signal. Above a level of signal noise all sharpness was lost, as anticipated. Thus the determining factors for boundary sharpness are curvature and separation speed, as the networks that maximise these two parameters produce the sharpest boundaries with or without signal noise (Fig. S18).

**Figure S18:**
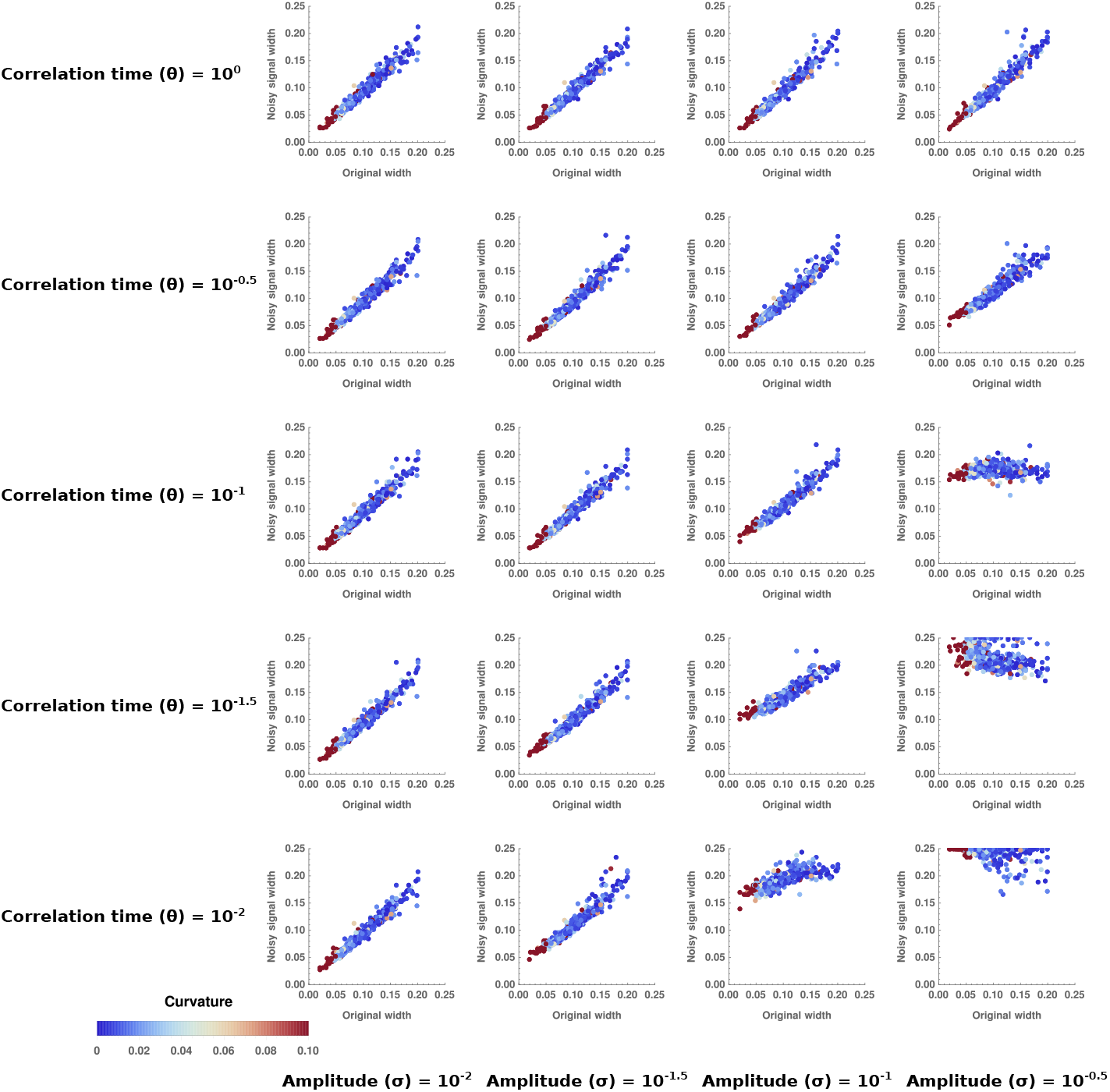
Effect of different levels of noise in the gradient on boundary precision. The boundary widths produced by systems recovered from the computational screen are plotted for simulations with no noise in the signal gradient (*x* axis) and with noise in the signal gradient (*y* axis). Colour labels networks from least (blue) to most (red) curvature. The behaviour of the noise in the signal has been modelled as an Ornstein-Uhlenbeck process, with the indicated amplitudes and correlation times. The same network parameter values were used for the simulations with and without signal noise. The analysis shows that the noise in the signal has relatively small effects on the precision of boundaries, except when the noise in the signal is so extreme that all sharpness is lost (bottom right plots).

### Comparison with Drosophila GAP gene and Eye Imaginal disc networks

We compared the networks recovered from the computational screen with those described for anterior posterior patterning of the Drosophila embryo and eye imaginal disc [Verd et al., 2017, Graham et al., 2010]. Both these systems have been characterised extensively such that we have sufficient knowledge of the network to perform our analysis [Akam, 1987, Ingham, 1988, Sánchez and Thieffry, 2001, Manu et al., 2009, O’Neill et al., 1994, Rebay and Rubin, 1995]. We added intrinsic noise to the original models from [Verd et al., 2017, Graham et al., 2010] using Langevin equations and an Ω that was chosen to result in fluctuations without leading to ergodicity. For the GAP gene system we used the parameters and equations as described in [Verd et al., 2017], for the imaginal disk network we used the Mathematica code provided as supplementary in [Graham et al., 2010]. We inspected the configurations in gene expression fluctuations near relevant steady states.

**Figure S19:**
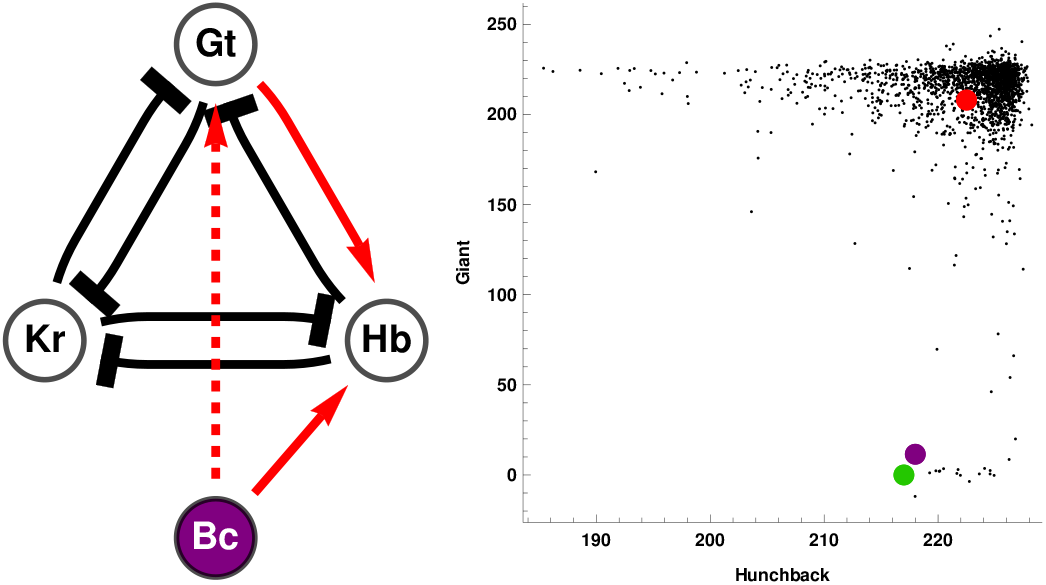
(Left) GAP gene network for the anterior boundary between Kruppel and Giant. The network has the same topology as the most frequently recovered network identified by our screen, with a difference only in the input (dashed link). This differences does not affect the dynamics as it only alters how the network interprets a change in signal. (Right) Black dots represent multiple simulations and illustrate fluctuations near the high-Giant / low-Kruppel steady state (red point). To transition to the low-Giant and high-Kruppel steady state (green point) the system must reach the transition point (purple point). The fluctuations in gene expression space are coerced into the Hunchback dimension (*x* axis), decreasing the probability of a stochastic fluctuation of the system reaching the transition point.

The architecture of the transcription circuits that comprise the GAP gene network [Verd et al., 2017] match closely those found in our computational screen (Fig. S16). As predicted by our computational screen, we can identify role for specific links between network components in the formation of GAP gene boundaries. In particular, for the anterior boundary between Giant and Kruppel, if we remove Knirps, which is not expressed in either of these domains, we find one the most common topologies recovered from our screen, with the correspondence Kruppel – *x*_1_, Hunchback – *x*_2_, Giant – *x*_3_ (Fig. S19). In this case, Hunchback and Giant display mutual exclusivity and the graded expression profile of Giant suggests that separation speed is used to sustain the sharp boundary; this is similar to the role played by Pax6 (*x*_3_) in the neural tube GRN. An interesting difference is that while Hunchback affects the direction of fluctuations in gene expression space, it does not change in concentration and simply alters the dynamics of the transition (Fig. S19).

**Figure S20:**
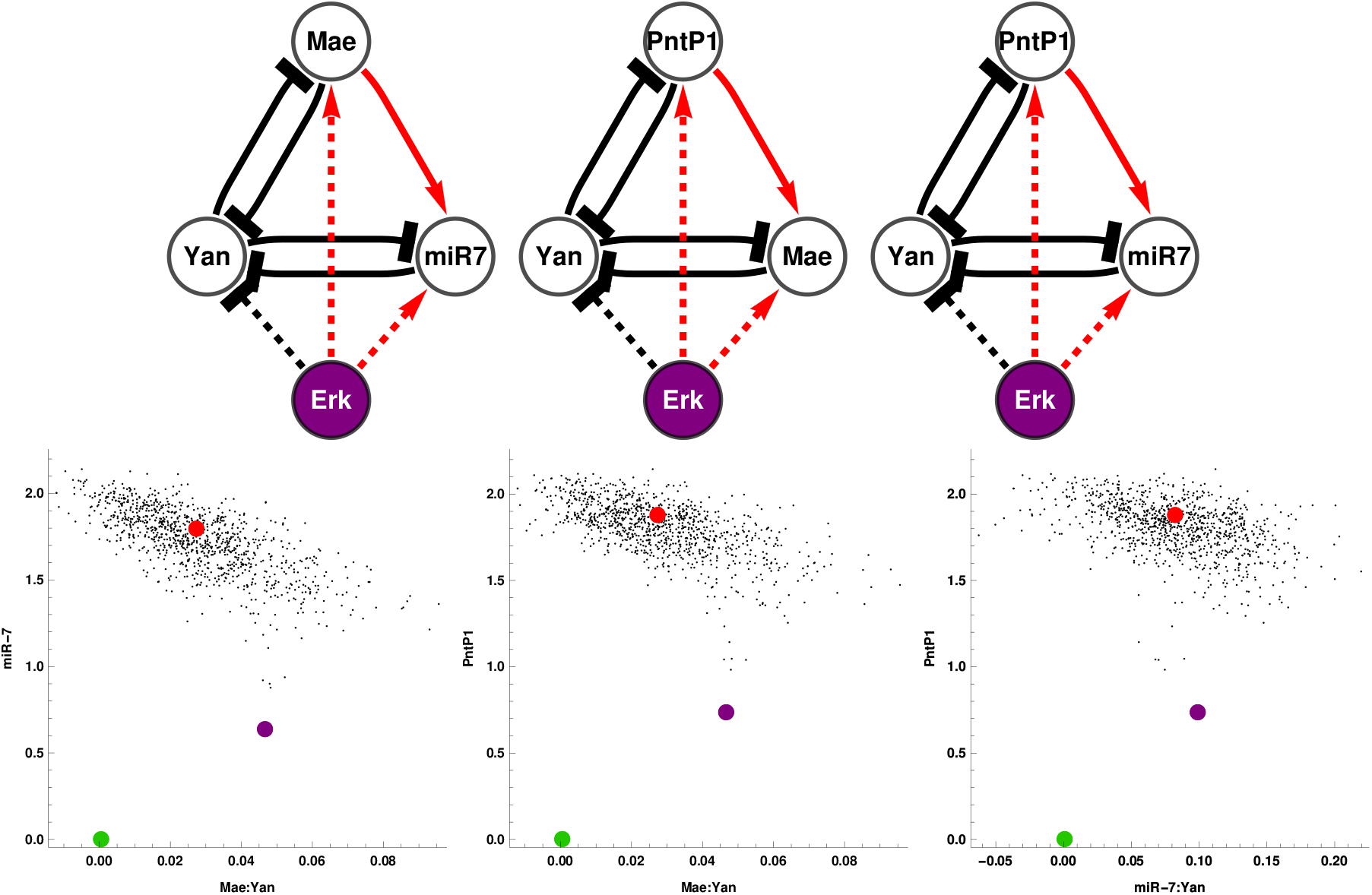
(Top) Multiple instances of one of the top topologies for boundary sharpness found in our computational screen are contained in the Drosophila eye disc network. This network is composed of several interactions and mediates the transition between Yan-on and Yan-off states. The configuration of inputs from the signal (Erk) is different to our topologies (dashed lines) but this does not affect the dynamics. (Bottom) Fluctuations near the Yan-off state (red point) for different projected views corresponding to the networks shown. The fluctuations are configured in directions that are not aligned with the transition point (purple point). This configuration decreases the possiblity of a cell reverting to a Yan-on state after the wave of Erk has shifted the system to a Yan-off state. Note that for Mae and miR-7 the inhibitions of Yan happen through direct interactions, thus where noted we show the fluctuations for the variable tracking the Inhibitor:Yan complex (Mae:Yan or miR-7:Yan).

The differentiation pattern of the eye imaginal disc also relies on cross-repressive interactions [O’Neill et al., 1994, Rebay and Rubin, 1995, Graham et al., 2010]. The expression of Yan, downstream of RTK signalling distinguishes between differentiated and undifferentiated precursors in the eye disc as the furrow migrates. We inspected the network proposed to achieve this [Graham et al., 2010] by focusing on three node networks that involved Yan and two other components in cross-repression with Yan. This approach resulted in three versions of a network topology found frequently as one with high curvature in our screen, with the mappings: (1) Yan – *x*_1_, miR-7 – *x*_2_, Mae - *x*_3_, (2) Yan – *x*_1_, Mae – *x*_2_, PntP1 – *x*_3_ and (3) Yan – *x*_1_, miR-7 – *x*_2_, PntP1 – *x*_3_ (Fig. S20). Simulations also indicate that the dynamics of these networks configure gene expression fluctations to decrease the probablity of a noise driven transition (Fig. S20). The bistable network facilitates a sharp switch between steady states, ensuring that cells only transition from a Yan-off to a Yan-on state when Yan is sufficiently activated by Erk signalling. Once the wave of Erk has passed, the dynamical curvature established by the network ensures that cells do not transition back to a Yan-on state (Fig. S20). Thus both Drosophila embryo and eye imaginal disc networks appear to have adopted network structures that are compatible with precision enhancing mechanisms.

## G Materials and methods

### G.1 Mouse Strains

Mouse strains containing the following alleles were used: Pax6(*Sey*) [Ericson et al., 1997] and O2e33^−/−^ in strain backgrounds C57BL/6Jax and F1(B6xCBA) respectively. The O2e33^−/−^ allele was derived using zygote injection of CRISPR gRNA and Cas9 plasmids (see below). Embryos were transferred to psuedopregnant females and subsequent pups were genotyped. O2e33^−/−^ mice were maintained as a heterozygous population; the line was sub-viable with less than 2/40 homozygous offspring surviving. Embryos for analyses were collected at the indicated time points following a mating, with the day of plug detection designated e0.5. All animal procedures were carried out in accordance with the Animal (Scientific Procedures) Act 1986 under the Home Office project licence PPL80/2528 and PD415DD17.

### G.2 Embryonic Stem Cell Culture

For the enhancer deletion *in vitro*, mouse ES cells containing a fluorescent reporter cotranslated with Olig2 (Olig2::T2A-mKate2) [Sagner et al., 2018] were used. Mouse embryonic stem cells were maintained on mitotically inactivated fibroblasts (feeder cells) in ES medium with 1,000 U/ml LIF. Cells were differentiated to spinal cord neural progenitors as previously described [Gouti et al., 2014]. To initiate differentiation, ES cells were dissociated using 0.05% Trypsin (Gibco) and panned in ES medium on culture plates for 2x 15 minutes to remove feeder cells. ES cells were collected, spun down and re-suspended in N2B27 medium. 50,000 cells were plates on 35mm CellBIND dishes (Corning). Dishes had been coated with 0.1% gelatine in PBS before addition of 1.5ml of N2B27 with 10 ng/ml bFGF. After 48 hours medium was replaced with N2B27 + 10ng/ml bFGF + 5uM CHIR99021 (Axon). 24 hours later, at D3, medium was replaced with N2B27 + 100nm RA (Sigma) and 500nm SAG (Calbiochem), this was repeated every 24 hours.

### G.3 CRISPR/Cas9 targeting

For CRISPR/Cas9-mediated excision of the −33 kb enhancer, two pairs of short guide RNA (sgRNA) sequences were designed to target either side of the enhancer region. ZiFit online tool (http://zifit.partners.org/) was used to select guides that had the lowest number of potential off target sites. sgRNA sequences (ACTTTGTAAGCCGAGCC) and (GATAATCGC-CTCCCTCC were cloned into pX459 v2.0 (Addgene, [Ran et al., 2013]) and transfected into ES cells via nucleofection. This generated a cell line with a 995bp deletion (chr16: 91192464-91193458). Two separate clones were analysed to determine whether there was substantial clonal variation. A second line was generated with a larger deletion of approximately 3.3kb using sgRNA sequences (GTTTATGGCTCATCCCC and TCCAGGCTCCCATATCC). Cell lines with this larger deletion yielded the same results as the smaller deletion (data not shown). To generate the mouse line, plasmids encoding the sgRNAs for the 3.3kb deletion were injected into zygotes before being transferred to pseudo-pregnant females. The mouse line generated had a 3259 bp deletion (chr16: 91191295-91194570).

To assess Olig2 protein copy number, a transgenic cell line was constructed, Olig2-HA-SnapTag. Sequencing encoding an HA tagged SnapTag was placed at the C-terminus of the endogenous coding sequence for Olig2 via homologous recombination using CRISPR. The SnapTag sequence was extracted from the pSNAPf vector (N9183S, NEB) and inserted into a plasmid containing Olig2 [Sagner et al., 2018] and targeted as previously described.

### G.4 Protein Copy Number Quantification

The concentration of recombinant proteins (used as standards) was calculated from Coomassie staining (GelCode Blue Stain Reagent, Thermo scientific). Recombinant proteins used were Pax6 (Bioclone, PI-0099) Nkx2.2 (MyBioSource, MBS717917) and SnapTag (NEB, P9312S). A solution of 5 Îĳm SNAP-tag was labelled with Janelia Fluor JF549 (TOCRIS, 6147) SnapTag Ligand at 10 Îĳm (assembled in house) for 30 mins at 37ÂřC.

To determine Pax6 and Nkx2.2 average molecule number per cell, a WT HM1 mouse embryonic stem cell line was used [Doetschman et al., 1987]. Cells were lysed in RIPA buffer supplemented with protease inhibitors. The cell lysates were analysed by Western blot, with lysate from a known number of cells loaded per lane. The following antibodies were used: rabbit anti-Pax6 (Millipore AB2237, 1:2000), mouse anti-Nkx2.2 (DSHB 745A5, 1:50), donkey anti-mouse IRDye 800CW (Licor) and donkey anti-rabbit IRDye 680RD (Licor). Blots were scanned using an Odyssey Scanner (Licor).

We used the cell line Olig2-HA-SnapTag to determine protein copy number for Olig2. Cells for Olig2 and Nkx2.2 copy numbers were differentiated as described. For Pax6, cells were exposed to 100nm RA only from day 4 to induce a more dorsal spinal cord cell fate. One day prior to sample collection, the cells were incubated with Janelia Fluor JF549 SnapTag Ligand (assembled in house) directly in the media at 1 *μ*M overnight. Cells were lysed in RIPA buffer supplemented with protease inhibitors. A known number of cells were loaded per lane. Gels were scanned using Typhoon FLA 9500.

To determine the percentage of expressing cells, flow cytometry was carried out as described in the Flow Cytometry section.

### G.5 Flow Cytometry Analysis

Cells were dissociated using 0.05% Trypsin and collected in ES media. Cells were then washed in PBS and resuspensed in PBS containing live-cell Calcein Violet dye (Life Technologies). Control and O2e33^−/−^ cells were differentiated in parallel and analysed together. Control cells differentiated without SAG from day 4 were used to set population gates for mKate positive cells.

For protein quantifications, flow cytometry was used to determine percentage of cells expressing Olig2, Pax6 and Nkx2.2. Cells were labelled with either PE Mouse anti-Nkx2.2 (BD Pharmingen 564730, 1:20); AlexaFluor 647 mouse anti-Human Pax6 (BD Pharmingen 562249, 1:50); goat anti-Olig2 (R&D Systems AF2418, 1:800) then donkey anti-goat 405 (Biotium 20398, 1:500). Flow analysis was performed using a Becton Dickinson LSRII flow cytometer.

### G.6 qPCR assays

The mRNA was extracted using RNeasy Mini Kit (Qiagen) according to the manufacturerâĂŹs instructions. 1 *μ*g of RNA was used for reverse transcription reaction using SuperScript III (Invitrogen) with random hexamers. Platinum SYBR Green qPCR SuperMix-UDG with ROX (Invitrogen) was used for amplification on a QuantStudio 5 Real-Time PCR system (ThermoFisher ScientiïňĄc). Expression values were normalised against *β*-actin. Two repeats of four (Islet1) samples at each timepoint were analysed. qPCR primers used were Islet1 FWD: 5âĂŹ-TATCAGGTTGTACGGGATCAAA and REV:5âĂŹ-CTACACAGCGGAAACACTCG.

### G.7 Immunohistochemistry and Microscopy

Embryos were collected at defined timepoints and fixed for 30 minutes for e8.5, 1 hour for e9.5 and 2 hours for e10.5 in 4% paraformaldehyde in PBS. Embryos for wholemount imaging were washed in PBS containing 0.1% Triton X-100 (PBST) before addition of primary antibodies. Embryos for sectioning were placed in cryopreservation 30% sucrose overnight at 4ÂřC then dissected into forelimb neural tube fragments. These were mounted in gelatine then frozen. 12Îĳm sections were collected on glass slides using Zeiss Hyrax C 60R cryostat. Gelatine was removed from the slides by 4 x 5 min washes in PBS at 42ÂřC and sections washed with PBST. For *in vitro* stainings, cells were washed in PBS and fixed in 4% paraformaldehyde for 15 min at 4ÂřC then washed in PBS then PBST. For whole embryos, embryo sections and cells, primary antibodies diluted in blocking solution (1% BSA in PBST) were applied overnight at 4ÂřC. These were then washed in 3 x PBST before secondary antibodies diluted in PBST were added for 1 hour at room temperature. Secondary antibodies were removed with 3 x washes with PBST and one wash containing PBST and DAPI. Sections and cells were mounted using Prolong Gold (Invitrogen). Embryos for wholemount were mounted using glycerol. Primary antibodies used were guinea pig anti-Olig2 (gift from Bennett Novitch, 1:8000 [Novitch et al., 2001]); mouse anti-Nkx2.2 (BD Pharmingen 564731, 1:500); rabbit anti-Pax6 (Millipore AB2237, 1:1000); goat anti-Sox2 (R&D Systems AF2018, 1:200); mouse anti-Mnx1/HB9 (DSHB 81.5C10, 1:40); rabbit anti-Olig2 (Millipore AB9610, 1:1000); goat anti-ISL1 (R&D Systems AF1837, 1:1000); mouse anti-Chx10 (Santa Cruz, sc-365519, 1:100). All secondary antibodies were raised in donkey and conjugated to Alexa488, Alexa568, Alexa647 (Abcam).

Cells were imaged on a Zeiss Imager.Z2 microscope using 20x objective. Z-stacks were taken and presented as a maximum projection using FiJi imaging software. A Leica SP5 upright confocal microscope was used to image embryo sections (40x oil objective) and whole embryos (20x dry objective). For Fig.2I, images were acquired using a Leica Sp8 inverted confocal (20x dry objective). For whole embryos, z-stacks were taken across a tile-scan then assembled and maximally projected using FiJi imaging software.

### G.8 Image quantification

#### Fluorescent intensity measurements

Single optical planes from confocal z-stack images were used for analysis. Each nucleus was identified individually using the FiJi point tool. The DAPI channel was used as reference for the position of the nuclei regardless of TF expression. A circle of 2 *μ*m radius was taken around each point, *x* and *y* position and mean fluorescence intensity values for Nkx2.2, Olig2 and Pax6 were recorded. Reference points at the ventral and dorsal pole of the neural tube in each section were recorded in order to align all embryos along the dorso-ventral axis.

#### Pre-processing

We performed a set of normalisation steps in order to compare embryos from different batches and across phenotypes:

1. The datasets were realigned vertically with respect to the reference points and the ventral-most point was set to (0,0) in axes coordinates
2. Cells with DAPI levels below two SDs from the mean were removed to eliminate falsely identified nuclei. This value was decided individually for each sample to account for different background levels resulting from technical noise.
3. Points that were very low in intensity (below two SDs) were set to a minimum threshold in each individual channel.
4. For Nkx2.2 and Olig2, the intensity values were re-scaled such that the minimum value is at 0 and the 40% quantile is at the arbitrary value of 0.08. This was done individually for each embryo with the assumption that most nuclei in a full neural tube cross-section will not express these proteins.
5. For Pax6, most nuclei in the image express some level of Pax6; accordingly we set the 60% quantile at 0.6 across all datasets.

#### Staging embryos with size

We used the dorsal-ventral length of the neural tube as a proxy for embryo age [Cohen et al., 2015]. For e9.5 embryos, the neural tube size measured was between 250*μ*m and 350*μ*m and for e10.5 embryos it was larger than 350*μ*m. In order to subgroup e9.5 embryos, neural tube size was used. In total we have 46 WT, 29 O2e33^−/−^ and 16 Pax6^−/−^. By sizes they are distributed as:

|  | WT | O2e33<sup>-/-</sup> | Pax6<sup>-/-</sup> |
| --- | --- | --- | --- |
| 150 - 250 $\mu\text{m}$ | 17 | 5 | 5 |
| 250 - 350 $\mu\text{m}$ | 13 | 13 | 3 |
| 350 - $\mu\text{m}$ | 16 | 11 | 8 |

#### Classification into cell types

In order to analyse the heterogeneity at the boundary between domains, we classified all cells into one of 5 specific cell types: floor plate, p3, pMN, Irx3 positive, other; this was done based on the position and expression profile of each cell. We refrained from using the Pax6 channel in our classifier to avoid any bias in the classification of Pax6^−/−^ embryos. We therefore classified based on three parameters: Nkx2.2 intensity, Olig2 intensity and dorsal-ventral position. The thresholds we employed for Nkx2.2 and Olig2 concentrations are shown in Fig. S21**A**-**B**. There was a further constraint on the dorsal-ventral position for each cell type, in order to avoid anomalies from blood vessels and imaging artefacts and to be able to separate floor plate cells from Irx3 positive cells, both of which lack expression of Nkx2.2 and Olig2 (Fig. S21**B**-**C**). Manually bench-marking this method indicated that we were able to classify most cells accurately for all three phenotypes. The classifier becomes less accurate for cells in dorsal regions but this is of no concern as our subsequent analysis did not involve these cells. For the specific task of quantifying the Olig2-Irx3 boundary position we employed the Pax6 channel as a further parameter to aid classification. This was only performed for WT and O2e33^−/−^ (data not shown).

**Figure S21:**
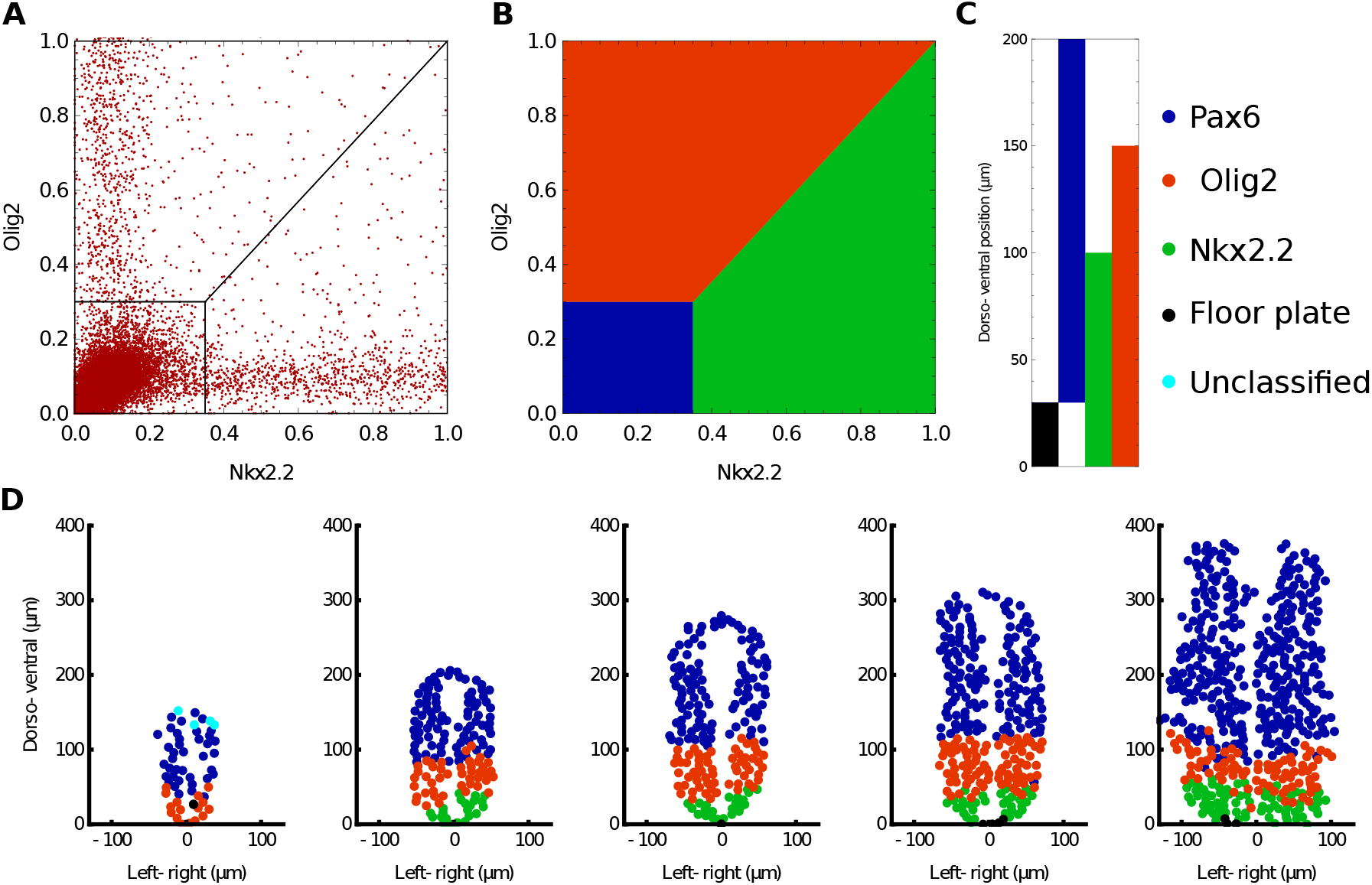
Analysis of gene expression in embryos. (**A**) Plot illustrating the concentrations of Nkx2.2 and Olig2 for all cells analysed. This highlights that the majority of cells are negative for both TFs and also that very few cells co-express both TFs. (**B**) Criteria to determine the identity of each cell by using the levels of Nkx2.2 and Olig2; colours indicating cell assignment as Olig2 (red), Nkx2.2 (green) and neither (blue) are consistent throughout the figure. The concentration of Pax6 is not used for classification. (**C**) Positional limits along the neural tube for each cell type. Cells that express neither Olig2 nor Nkx2.2 are classified based on their position as they can be ventral floor plate cells (black) or more dorsal progenitors. Cells that have mismatching values of concentration and position are classified as exceptions in Cyan (**D**) Examples of classified embryos of increasing age, illustrating the accuracy of the approach for determining cell type.

#### Defining boundary position and width

Once the cell types had been classified we assigned a quantitative measure of the width of gene expression boundaries. For this we fit to the cell position data, for each embryo, a smooth function indicating the probability of finding a cell of one type (the prevalent type on one side of the boundary) at each location of the image. We focused on the boundary between p3 and pMN domains. The classifier is then binary and gives the probability of finding a p3 cell at each image location. We used a Gaussian process approach to fit this classifier as detailed in [Rasmussen and Williams, 2004], using public MATLAB code (MATLAB version r2018b). The Gaussian process was chosen to have a constant mean function and a squared exponential covariance function. This choice of covariance function is relatively standard and allows us in particular to assign separate covariance function lengthscales in the *x* and *y* image directions by automatic relevance determination [Rasmussen and Williams, 2004]. We used a logistic transfer function to convert Gaussian process values to probabilities, again a standard choice. Once the classification probabilities have been obtained in this way, we define the boundary as the region where the probability of p3 cells lies in the range 11% to 89%, i.e. where there is significant mixing of cell types. We then determine the width of this region geometrically. This method allowed us to calculate the boundary widths for all embryos in a consistent manner, and to compare WT with mutants. The boundary region is determined from the trained classifier for each embryo as explained above; the position where the classification probability is 50% for either cell type is used to define the position of the boundary (an average position of the boundary along the left-right axis) (Fig. S22). We do not use entropy based measures such as in [Dubuis et al., 2013, Petkova et al., 2019] as these typically rely on the assumption of Gaussian gene expression level distributions; this assumption is inapplicable in the boundary region, where the system is bistable and the distributions therefore bimodal. Information theoretical methods are also normally used with a single spatial coordinate while we are looking at a 2D tissue. This may have irregular growth or oblique sectioning which could lead to a slanted boundary and therefore misleading results once projected onto a single dimension.

#### Quantifying TF levels

We extracted Olig2 positive cells that were classified as being within the boundary region. The model predicted that these cells were the most likely to transition to a Nkx2.2 positive state, given sufficient time. We quantify the levels of Pax6 and Olig2 for these cells in WT and O2e33^−/−^ mutants. The resulting measurements do not provide absolute numbers; but given that all samples are normalised in the same way, as described (Sec. G.8), the resulting measurements are comparable relative to each other. We use these measurements as equivalents to observing fluctuations around a steady state over a series of dorso-ventral positions. In this way, we take the corresponding equivalent in the simulations, where we also average fluctuations across several neural tube positions (Supp. C).

**Figure S22:**
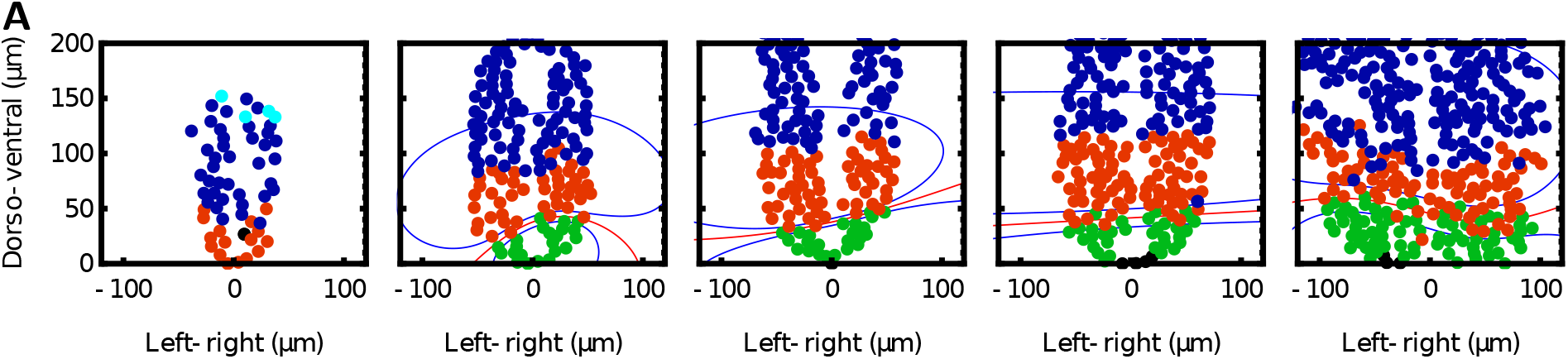
Examples of boundaries determined by the Gaussian process classifier. The red lines indicate the computed boundary position, and correspond to the image locations where the probability of being a p3 or pMN cell is 0.5. Blue lines close to the p3-pMN boundary delimit the area identified as the boundary region, where the probability of being a p3 cell is in the range 11% to 89%. By measuring the area between the two blue curves and dividing by the width of the embryo we are able to quantify the width of the boundaries. In turn by obtaining the average position of the red line, we are able to calculate the boundary position.

#### Calculating variance levels

In order to calculate the total variance of Olig2 and Pax6 levels within the pMN domain we extracted all Olig2 expressing cells, for both WT and O2e33^−/−^, outside the boundary region. The variances and covariances of the normalised fluorescence intensity values were calculated, in analogy with the theoretical approach (Supp. C). The square root of the trace of the resulting covariance matrices was then used to obtain the typical root-mean-square relative variance.

